# Repurposing the plant-derived compound apigenin for senomorphic effect in antiaging pipelines

**DOI:** 10.1101/2024.09.09.611999

**Authors:** Hongwei Zhang, Qixia Xu, Zhirui Jiang, Rong Sun, Sanhong Liu, James L. kirkland, Weidong Zhang, Yu Sun

**Author notes:** Co-correspondence: Weidong Zhang, Yu Sun. E-mail addresses (W.Z.); (Y.S.). These authors contributed equally: Hongwei Zhang, Qixia Xu.

## Abstract

Cellular senescence is a cell fate triggered by inherent or environmental stress and characterized by stable cell cycle arrest accompanied by a hypersecretory feature, termed as the senescence-associated secretory phenotype (SASP). Senescent cell burden increases with natural aging, functionally contributing to age-related organ dysfunction and multiple disorders. In this study, we performed a large scale screening of a natural product library for senotherapeutic candidates by assessing their effects on human senescent cells. Apigenin, a dietary flavonoid previously reported with antioxidant and anti-inflammatory activities, exhibited a prominent capacity in targeting senescent cells as a senomorphic agent. In senescent cells, apigenin blocks the interactions between ATM/p38 and HSPA8, thus preventing transition of the acute stress-associated phenotype (ASAP) towards the SASP. Mechanistically, apigenin targets peroxiredoxin 6 (PRDX6), an intracellular redox-active molecule, suppressing the iPLA2 activity of PRDX6 and disrupting downstream reactions underlying the SASP development. Without reversing cellular senescence, apigenin deprives cancer cells of malignancy acquired from senescent stromal cells in culture, while reducing chemoresistance upon combination with chemotherapy in anticancer regimens. In preclinical trials, apigenin administration improves physical function of animals prematurely aged after whole body irradiation, alleviating physical frailty and cognitive impairment. Overall, our study demonstrates the potential of exploiting a naturally derived compound with senomorphic capacity to achieve geroprotective effects by modulating the SASP, thus providing a research platform for future exploration of novel natural agents against age-related conditions.

## INTRODUCTION

Demographic data of recent years increasingly show a global pattern of human aging, while chronic age-related diseases in the elderly represent an enormous burden and unprecedented challenge to medical healthcare ^1^. Cellular senescence is one of the central hallmarks of aging, also an emerging therapeutic target exploitable to intervene the aging process *per se* and related disorders ^2, 3^. Although senescence appears to be beneficial in a few physiological settings such as embryonic development, tissue repair and wound healing, increasing lines of evidence suggest that most, if not all, human pathologies can be promoted or caused by the accumulation of senescent cells in tissues and organs ^4, 5^. Recent preclinical studies have intensively explored the therapeutic potential of targeting senescence-related pathways in aging and diseases, while more investigations particularly those involving future clinical trials, are essential to understand the efficacy and safety of senotherapy, a novel therapeutic modality that minimizes, or at least, ameliorates the detrimental effects of senescent cell accumulation with aging ^6^.

Senotherapeutics, a group of pharmacological agents that specifically target senescence, typically include those attenuating the pro-inflammatory activity of senescent cells (senomorphics) and those inducing preferential lysis of senescent cells (senolytics) ^7, 8^. Elimination of senescent cells markedly increases mobility, improves physical condition, enhances cognitive function and extends overall lifespan in aged or diseased animals ^9–11^. Notably, administration of senolytics also confers additional health benefits in human patients, as evidenced by the case of dasatinib and quercetin (‘D + Q’) combination in clinical trials for diabetic kidney disease (DKD) and idiopathic pulmonary fibrosis (IPF) ^12–14^]. Initial outcomes from a vanguard clinical trial conducted in early-stage symptomatic patients with Alzheimer’s disease (AD) aimed to assess central nervous system (CNS) penetrance further supported the safety, feasibility and efficacy of these agents and provided mechanistic insights of senolytic effects ^15^. In contrast to senolysis, attenuating the influence of senescent cells *via* senomorphic agents represents another effective approach ^4, 16^. An ideal senomorphic candidate should be able to target senescent cells by blocking development of the senescence-associated secretory phenotype (SASP), rather than killing senescent cells or eliciting their re-entry to cell cycle, since rejuvenated cells may still carry a mutation load ^17^. Phytochemicals, including alkaloids and polyphenols, have demonstrated the ability to target senescent cells, either reducing their burden *in vivo* and/or modulate their SASP, together enhancing the potential of organ rejuvenation ^18, 19^. Interestingly, some natural agents exhibit both senomorphic and senolytic activities, largely depending on the dose to be used and/or pathological settings to intervene ^11, 20^.

The drug development pipeline has been inefficient, as only 15.3% agents in phase 1 clinical trials can advance to FDA approval in the US ^21^. Repurposing drugs formerly approved for clinical use represents an efficient tactic to circumvent drug development pitfalls and to reduce overall costs. Panels of small molecules for drug repurposing studies can be procured and customized, providing an opportunity to adapt drug repurposing screens for alternative indications. As a naturally available flavonoid in fruits and vegetables, apigenin has been reported with multiple biological activities including antioxidant, anti-inflammatory, anti-viral and anticancer properties ^22^. By reducing reactive oxygen species (ROS) production while enhancing glutathione and mitochondrial adenosine triphosphate levels, apigenin effectively prevents abnormal mitochondrial distribution and early apoptosis in oocytes, thus minimizing the deterioration of oocyte quality during aging ^23^. It can significantly reverse the glutathione reduction tendency while decreasing malondialdehyde and catalase activity in the brain, displaying potential anti-depressant effects in animal models ^24^. Apigenin can suppress the SASP partially by inhibiting IL-1α signaling *via* IRAK1, IRAK4, p38 MAPK and NF-κB, resulting in declined aggressiveness of human breast cancer cells ^25^. However, the benefit of apigenin in a wider range of age-related settings, particularly those involving cellular senescence in the advanced life stage and its potential value in antagonizing pathologies and physical dysfunction, remains yet underexplored. In this study, we screened a library of natural medicinal agents (NMAs) for potential effects to modulate the survival and pro-inflammatory activity of human primary cells induced senescent *in vitro*. Our results support the competency of apigenin in targeting human senescent cells as a novel senomorphic agent. We further unmasked its functional mechanism and pharmacological value in restraining senescence-associated activities as exemplified by the case of tumor progression and premature aging. This study demonstrates the benefits of apigenin, a natural compound, in curtailing the pathological influence of senescent cells, thus providing a baseline for its future development and application in geriatric medicine.

## 2. RESULTS

### 2.1 Drug screening discloses the potential of apigenin as a senomorphic agent

In an effort to discover novel natural agents that can effectively regulate senescent cells, we performed a comprehensive drug screening with a NMA library composed of 66 natural products (Table S1), mostly secondary metabolites derived from plants, microbes and certain animals. To this end, PSC27, a primary normal human prostate stromal cell line, was selected as an experimental cell model for extensive studies. PSC27 develops a canonical SASP upon senescence induced by inherent stress and/or exterior stimuli, including exhaustive replication (replicative senescence, RS), genotoxic chemotherapy and ionizing radiation (therapy-induced senescence, TIS) ^26–29^. We exposed cells to a sub-lethal dose of bleomycin (BLEO, 50 μg/ml), with a typical cellular senescence phenotype observed 8-10 d after treatment, as evidenced by elevated senescence-associated β-galactosidase (SA-β-Gal) staining positivity, decreased BrdU incorporation and increased DNA-damage repair (DDR) foci (Figure S1a-c). To streamline, we designed a screening procedure to evaluate the effects individual natural agents exerted on the *in vitro* feasibility of senescent cells and expression of their SASP (Figure 1a).

**Fig. 1.**
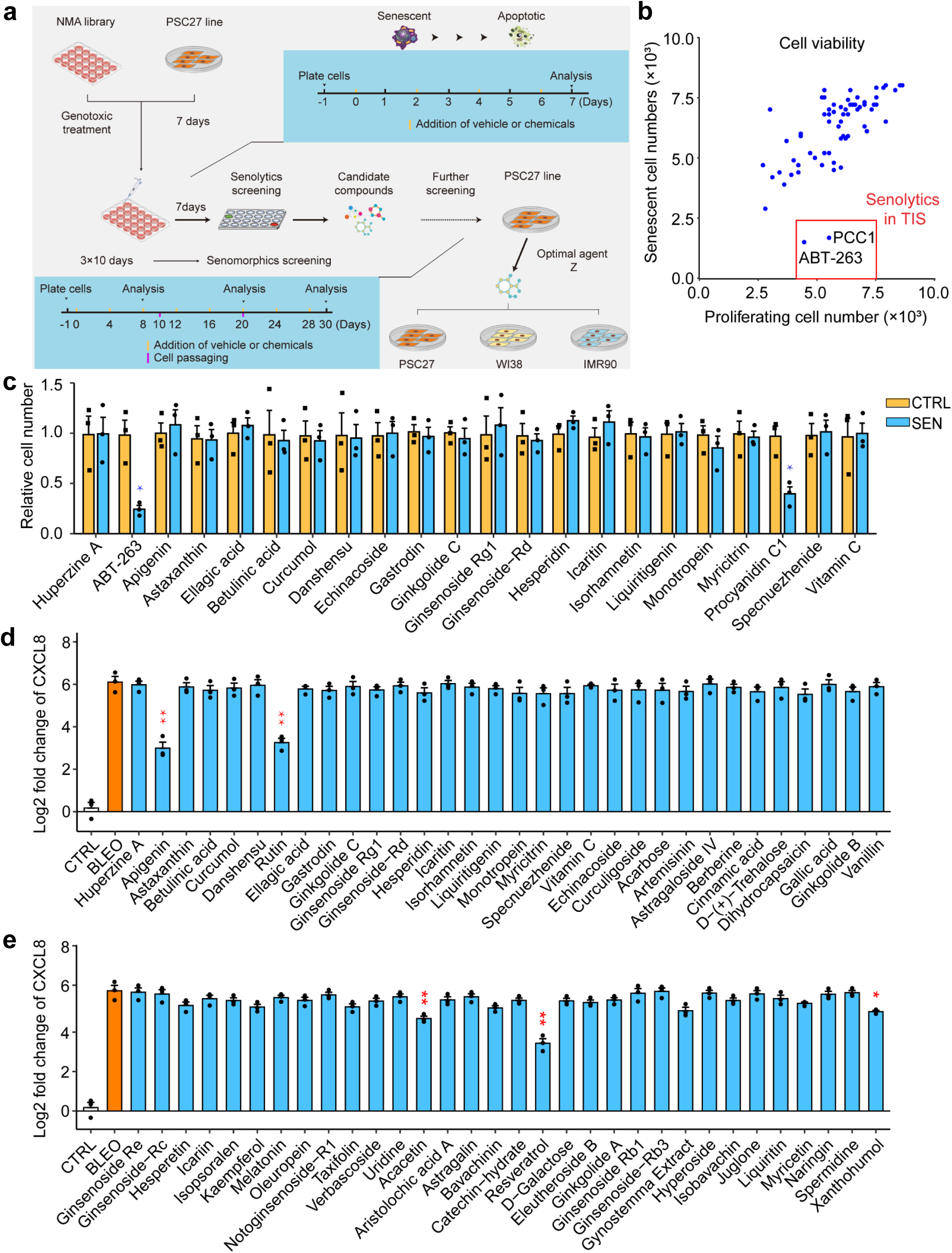
*In vitro* screening of a NMA library for potential senotherapeutics. (a) A schematic representation of the procedure for human cell-based screening of a NMA library composed of 66 naturally-derived agents. Screening was first performed for all natural compounds for senomorphics, with the efficacy of potential candidates further validated in multiple human stromal cell lines including PSC27, WI38 and IMR90. (b) Experimental outputs of senolytics screening after pharmacological treatments *in vitro*. Natural compounds were assayed by incubation for 3 days at a concentration of 3 μg/mL/agent (ABT-263 at 1.25 μM) with 5.0×10^3^ cells. Each hit denoted by a blue dot is shown based on the effect to selectively kill SEN but not CTRL cells, and represents the mean of 3 biological replicates. The red-edged rectangular region shows candidates of senolytics. Note, only ABT-263 and procyanidin C1 (PCC1), the positive controls, fell in this specific region. (c) Evaluation of the efficacy of natural compounds as senolytics in SEN and CTRL counterpart cells. (d-e) Assessment of the efficacy of natural compounds (d, group A; e, group B) by analyzing the expression of a canonical SASP soluble factor CXCL8. For all datasets, samples were examined after treatment with individual agents in culture for 3 days. NMA, natural medicinal agent. CTRL, control. SEN, senescent. TIS, therapy-induced senescence. Data in b, c, d and e are shown as mean ± SD and representative of 3 independent biological replicates. *P* values were calculated by Student’s *t* tests. ^, *P* > 0.05; *, *P* < 0.05; **, *P* < 0.01; ***, *P* < 0.001; ****, *P* < 0.0001.

Induction of apoptosis in senescent cells represents a central feature and the most valuable capacity of senolytics, as exemplified by studies involving senolytic compounds such as ABT-263, ABT-737 and procyanidin C1 (PCC1) ^28, 30, 31^. Alternatively, a major advantage of senomorphics, for example, rutin and resveratrol ^29, 32^, is the ability to selectively downregulate the SASP expression. To determine the relevant anti-senescence potentials in these NMA components, we analyzed the effect of these natural agents on senescent PSC27 cells, by first confirming its experimental feasibility as a cell-based model for geroprotective compound screening purpose. Preliminary results indicated that some agents either killed senescent cells (while preserving their proliferating counterparts), or suppressed the expression of CXCL8, a canonical SASP factor, thus further validating the potential and appropriateness of PSC27 for extended investigations (Figure 1b-e and Figure S1d,e). Despite a thorough screening of the NMA library, we failed to discover any new senolytic agents, which are supposed to functionally resemble the positive controls ABT-263 and PCC1 (Figure 1b,c and Figure S1d,e). The result also implied the rareness of natural senolytic agents and the challenge in expanding the present reservoir of senotherapeutic agents particularly those of senolytic potential. Nevertheless, we noticed that several natural compounds generated a significant senomorphic effect, which warrants in-depth studies (Figure 1d,e).

Among the medicinal agents, rutin, resveratrol and apigenin displayed a strong senomorphic capacity (Figure 1d,e). Resveratrol is natural product with SASP-inhibiting activity and targets PI3K/Akt signaling pathway ^32^. As another phytochemical molecule recently unraveled with senomorphic efficacy, rutin attenuates the acute stress-associated phenotype (ASAP) and affects the SASP development afterwards, mainly through disrupting the interactions between ATM and HIF-1α as well as between ATM and TRAF6 ^29^. In this study, we chose to specifically focus on apigenin, a plant-derived natural flavonoid, which can downregulate the expression of typical SASP factors upon bleomycin-induced senescence in BJ fibroblasts by inhibiting NF-κB p65 activity through targeting IRAK1/IκBα signaling and IκBζ expression, with an *in vivo* activity of SASP suppression identified in the kidney of aged rats ^33^. Apigenin can also restrain the SASP by blockade of IL-1α signaling through IRAK1, p38 MAPK and NF-κB, thus alleviating pro-tumorigenic effect of the SASP in breast tumors ^25^. However, despite these intracellular activities, which are relative downstream in the SASP signaling network, a wider influence of apigenin on the transcriptome-wide expression of senescent cells, especially the more upstream or even direct targets after exposure of these cells to apigenin, remains basically unknown.

### 2.2 Apigenin prominently attenuates the SASP but not cellular senescence

Given that apigenin inhibits the expression of CXCL8, a SASP hallmark factor, we further asked whether it downregulates the vast majority of SASP factors, or even counteracts cellular senescence. Therefore, we performed SA-β-Gal staining and BrdU incorporation assays *in vitro* to determine the anti-senescence effects of apigenin. The data indicated that SA-β-Gal staining profile and mitotic inactivity remained largely unchanged, in both proliferating cells and their senescent counterparts (Figure 2a,b).

**Fig. 2.**
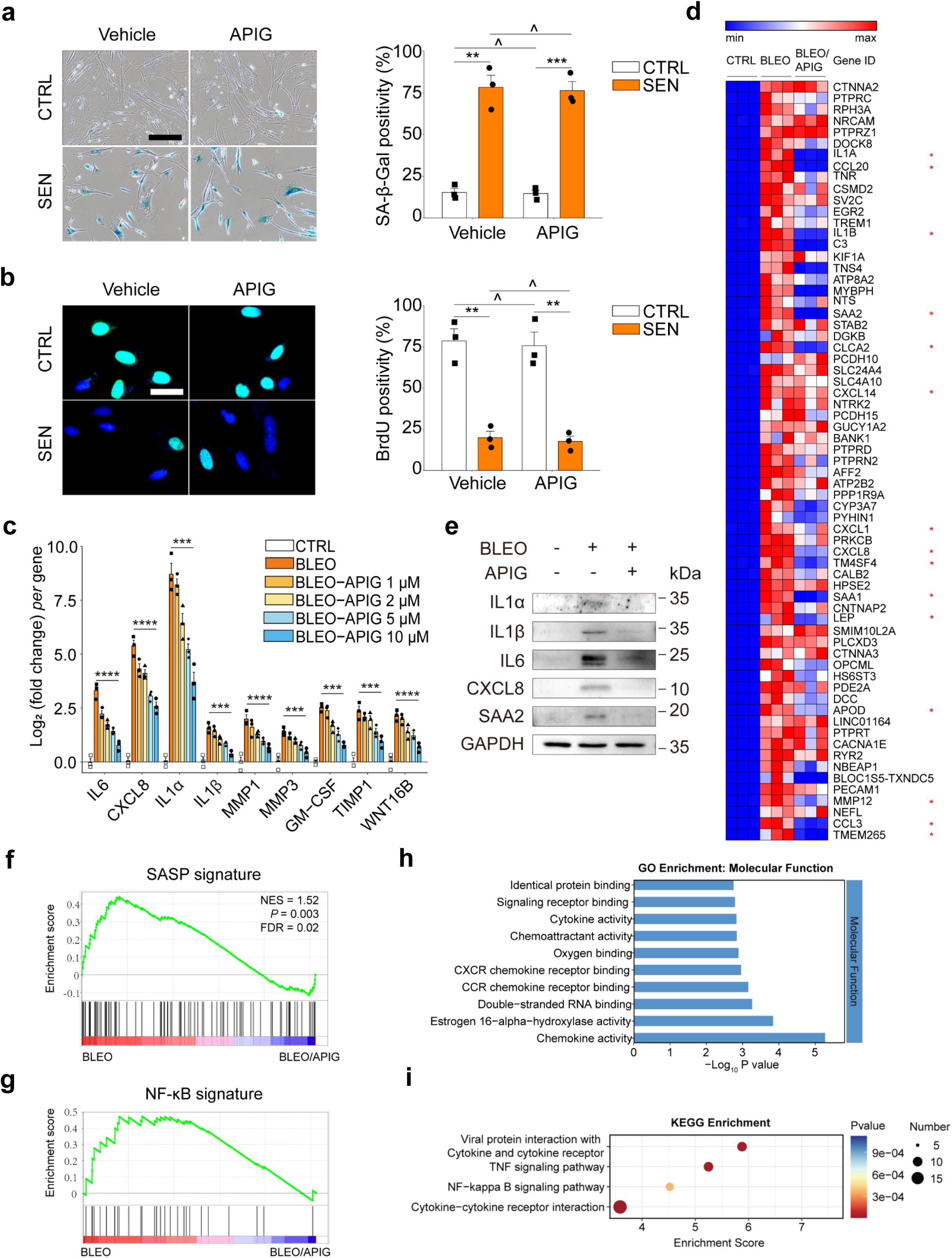
Characterization of the senomorphic potential of apigenin. (a) Senescence examination by SA-β-Gal staining of PSC27 cells. Left, representative images. Scale bar, 20 µm. Right, statistics. (b) DNA synthesis assessment by BrdU incorporation. Left, representative images, Scale bars, 5 µm. Right, statistics. (c) Quantitative analysis of the expression of canonical SASP factors at the transcription level upon BLEO-induced senescence and treatment by apigenin at increasing concentrations. (d) Hierarchical clustering heatmap depicting top human genes (67) that were significantly upregulated in senescent stromal cells but downregulated upon apigenin treatment (10 μM). Red stars, representative SASP factors. (e) Immunoblot analysis of typical SASP factors at protein levels of stromal cells exposed to BLEO and/or apigenin. GAPDH, loading control. (f) GSEA profiling of enrichment scores of typical soluble factors in the wide SASP spectrum. (g) GSEA profiling of enrichment scores of factors related to NF-κB pathway. (h) A bar chart shows the GO molecular function of differentially expressed genes (67) depicted in (d). (i) KEGG pathway enrichment analysis of differentially expressed genes (67) depicted in (d). Unless specially noted, data in a, b and c are shown as mean ± SD and representative of 3 independent biological replicates with *P* values calculated by Student’s *t* tests. ^, *P* > 0.05; *, *P* < 0.05; **, *P* < 0.01; ***, *P* < 0.001; ****, *P* < 0.0001.

A subset of hallmark SASP factors, including IL6, CXCL8, IL1α/1β, MMP1/3, GM-CSF, TIMP1 and WNT16B displayed a dose-dependent decline upon exposure of senescent cells to apigenin, with a concentration of 10 μM appeared most effective (Figure 2c). Furthermore, transcriptomic data from RNA-seq showed that expression of the canonical SASP factors was markedly upregulated upon cellular senescence but significantly downregulated upon apigenin treatment (Figure 2d). Immunoblot assays indicated that apigenin abrogated the SASP expression in senescent cells, as evidenced by the decreased expression of IL1α/1β/6, CXCL8 and SAA2 (Figure 2e). Given the striking effect of apigenin in restraining the SASP expression, we performed GSEA analysis for signatures of SASP and NF-κB, the latter plays a significant role in activating the ASAP and sustaining the SASP ^27^. Our results suggested that both molecular signatures were apparently suppressed upon apigenin treatment (Figure 2f,g).

Gene ontology (GO) enrichment analysis of apigenin-downregulated genes indicated that chemokine activity, estrogen 16-alpha-hydroxylase activity, double-stranded RNA binding, CCR/CXCR chemokine receptor binding activities were among the most significantly suppressed molecular functions (MF) (Figure 2h). In terms of biological processes (BP), type I interferon signaling pathway, defense response to virus, response to virus and inflammatory response were most dramatically affected (Figure S2a). The representative cellular components (CC) most associated with downregulated genes were those residing in extracellular region and space, although some proteins were transported to plasma membrane, platelet dense granule lumen, as well as calcium- and calmodulin-dependent protein kinase complex (Figure S2b). KEGG pathway analysis suggested that cytokine-cytokine receptor interaction, viral protein interaction with cytokine receptor, TNF signaling and NF-κB signaling pathways were most significantly inhibited upon treatment of senescent cells by apigenin (Figure 2i). Altogether, all these data substantiate a prominent capacity of apigenin in suppressing the expression of pro-inflammatory cytokines and chemokines, a subset of central SASP factors, starting from the transcriptomic level.

We next performed proteomics analysis with whole cell lysates of cells using mass spectrometry, an approach that allows to reveal senescence-related protein level changes and offer potential therapeutic targets ^34^. Largely resembling transcriptomic alterations, our proteomic data indicated that apigenin treatment caused a remarkable downregulation of senescence-related proteins, such as EIF4A2, CCND3 and ITPR3 (Figure S2c,d). Among them, EIF4A2 (eukaryotic translation initiation factor 4A2) is negatively correlated with mRNA translation upon oncogene-induced senescence (OIS) and can promote extracellular matrix deposition, thereby holding the capacity to shape the secretory protein translational landscape ^35^. CCND3 (cyclin D3) is an activator of cellular senescence and promotes the hypermitogenic arrest, a status that is stimulated by mitogens and defines cellular senescence as a functionally active, stable and conditionally reversible state ^36^. ITPR3 (inositol 1,4,5-trisphosphate receptor type 3) promotes cellular senescence by mitochondrial calcium accumulation, while its knockdown enhances OIS escape ^37, 38^.

To expand, we further assessed the effect of apigenin on senescent cells induced by other means, including inherent stress or environmental stimulation, as illustrated by replicative senescence (RS) and ionizing radiation (RAD), respectively. As expected, apigenin holds the potential to diminish the expression of the vast majority of canonical SASP factors, regardless of the types of senescence-inducing modality (Figure S2e,f). Additionally, we performed further experiments with WI38 and IMR90, two typical fibroblast lines derived from human embryonic lung tissues. The results largely resembled that of PSC27, a human prostate-derived stromal line, suggesting that apigenin-exerted suppression of the SASP is essentially organ type-independent (Figure S2g,h). Together, our studies demonstrated a salient capacity of apigenin in restraining the SASP expression, which is neither senescence type- nor organ type-dependent.

### 2.3 Apigenin dampens the SASP expression by interfering with interactions of ATM and p38 MAPK with HSPA8

Given the critical role of apigenin in suppressing the SASP, we next queried its functional mechanism in targeting senescent cells. To examine the potential effect of apigenin on DDR signaling, which is usually the driving force of a persistent SASP, we first investigated the extent of molecular pathway perturbation in senescent cells upon exposure to apigenin. Induction of senescence by BLEO caused a notable activation of ATM, a central modulator of the DDR signaling cascade, as evidenced by immunoblot assays (Figure 3a). As a response to external challenge or endogenous stimuli, proliferating cells first develop an acute stress-associated phenotype (ASAP), characterized by phosphorylation and nucleus-to-cytoplasm translocation of ATM, TRAF6 auto-ubiquitination and TAK1 activation, a series of prompt intracellular reactions to insulting damages before senescence markers become evident ^27^ (Figure 3a). Upon cell treatment by genotoxicity, TAK1, a key cytoplasmic kinase in modulation of the SASP, was rapidly phosphorylated, a change allowing its functional involvement in the transition of the ASAP towards the SASP via dual feedforward mechanisms ^27^. We further observed phosphorylation/activation of p38 MAPK, a molecule downstream of TAK1, followed by engagement of PI3K/Akt/mTOR, a signaling pathway that supports the formation of a chronic secretory phenotype, namely the SASP. However, in the presence of apigenin, these sequential molecular events were generally abrogated, although phosphorylation of ATM and TAK1 remained largely unaffected. According to such a molecular pattern, we would extrapolate that the potential target(s) of apigenin may be mechanistically located downstream of ATM and/or TAK1, but upstream of p38 MAPK, PI3K/Akt/mTOR and other factors, although with a possibility to be implicated in the transition of the ASAP to the SASP during cellular senescence.

**Fig. 3.**
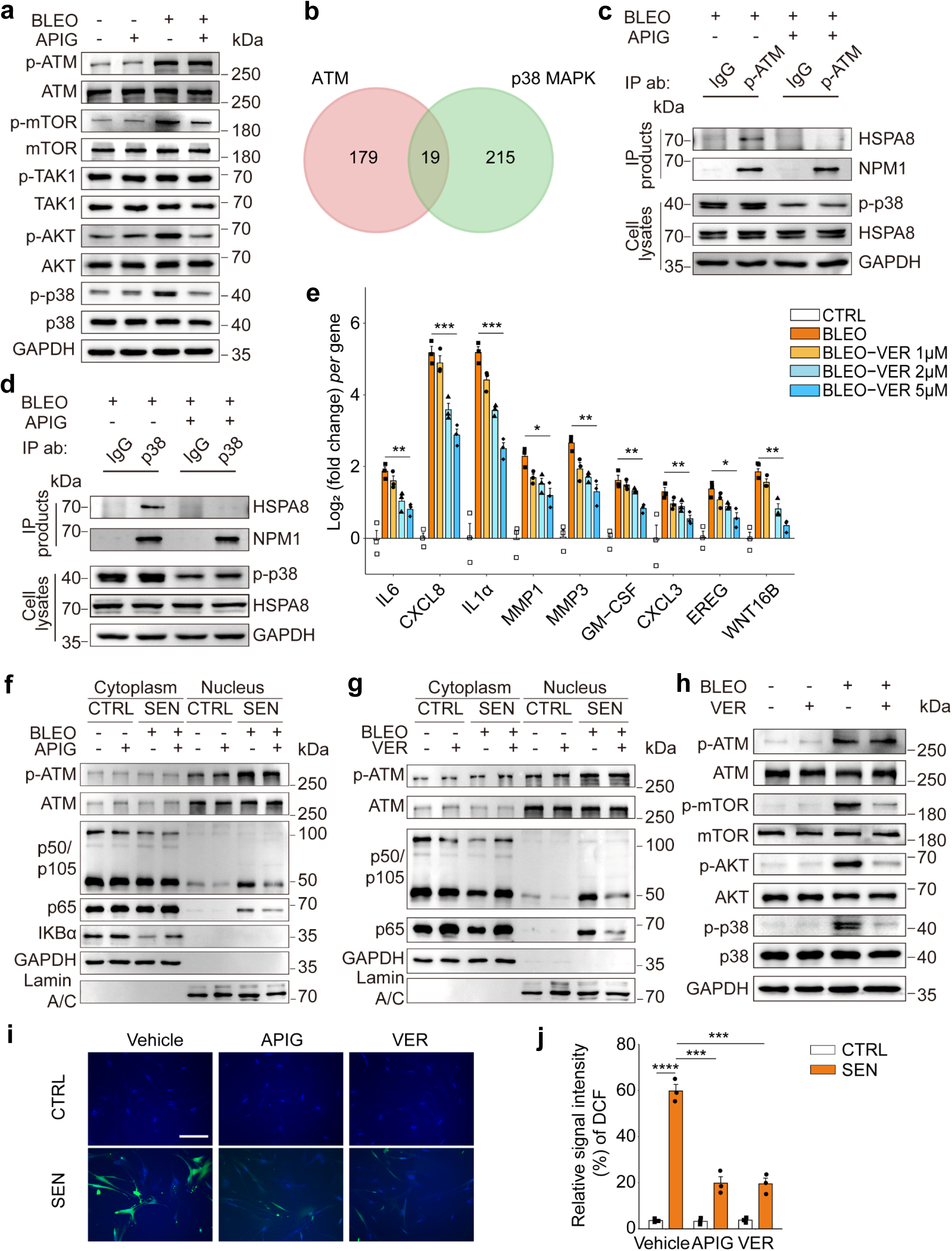
Apigenin restrains the SASP expression by interfering with DDR signal transduction in senescent cells. (a) Immunoblot analysis of DDR signaling-related molecules involved in regulation of the SASP expression in mammalian cells. After 7 d incubation with BLEO and/or apigenin, PSC27 cells were lysed for examination. GAPDH, loading control. (b) Venn plot depicting 19 mutually interacting molecules shared by both ATM and p38 MAPK uncovered by bioinformatics mining. (c) Immunoprecipitation (IP) assay coupled with immunoblot analysis to detect protein-protein interactions. PSC27 cells were treated with BLEO (50 μg/mL) for 12 h to induce senescence, in the absence of presence of apigenin in culture for 7 d. Cells were then lysed for IP with IgG or anti-p-ATM, with HSPA8 and NPM1 in both immunoprecipitates (IPs) and inputs examined. (d) IP coupled with immunoblot to probe protein-protein interactions. Sample preparation was performed as described in (c). Lysates were immunoprecipitated with IgG or anti-p38, with HSPA8 and NPM1 in both IPs and inputs examined. (e) Quantitative assessment of canonical SASP factor expression at transcriptional level upon senescence induced by BLEO at increasing concentrations of VER155008 (VER), a HSPA8 inhibitor. (f) Immunoblot analysis of ATM, p50/105, p65 and IKBα translocation between nucleus and cytoplasm upon treatment with apigenin. Lamin A/C and GAPDH, loading controls for nucleus and cytoplasm, respectively. (g) Immunoblot analysis of ATM, p50/105, p65 and IKBα translocation between nucleus and cytoplasm. Samples preparation was similar to that described in (f), except that VER155008 was applied instead of apigenin. Lamin A/C and GAPDH, loading controls for nuclei and cytoplasm, respectively. (h) Immunoblot assessment of DDR signaling-related molecules involved in regulation of the SASP expression. After 7 d incubation with BLEO with or without VER, PSC27 cells were lysed for analysis. GAPDH, loading control. (i) Measurement of ROS level *via* 2′-7′-dichlorodihydrofluorescein diacetate (DCFH-DA), a ROS-sensitive probe, in proliferating or senescent cells in the presence or absence of apigenin in culture 1 d afterwards. Representative images are shown. Scale bar, 20 μm. (j) Comparative statistics of ROS signals imaged by the DCFH-DA fluorescent probe. DDR, DNA damage response. BLEO, bleomycin. ROS, reactive oxygen species. Unless specially noted, data in e and j are shown as mean ± SD and representative of 3 independent biological replicates, with *P* values calculated by Student’s *t*-tests. ^, *P* > 0.05; ^, *P* > 0.05; *, *P* < 0.05; **, *P* < 0.01; ***, *P* < 0.001; ****, *P* < 0.0001.

To unravel the mechanism regulating the function of apigenin in targeting senescent cells, we conducted a proteome-scale profiling of interactors with a capacity to interact with both ATM and p38 MAPK, two molecules that are known to be involved in the transduction of signals to allow the SASP development but lack direct correlation in senescent cells. Bioinformatics suggested 198 ATM- and 234 p38 MAPK-unique interacting molecules in human cells, with 19 such interactors shared by both ATM and p38 MAPK (Figure 3b and Figure S3a,b). Among these interactor candidates, we noticed that HSPA8 and NPM1, intracellular molecules that hold the potential to mediate the interaction between ATM and p38 MAPK, thus deserving further investigation. The p-ATM-mediated co-immunoprecipitation (co-IP) revealed that each of HSPA8 and NPM1 can direct interact with p-ATM and p38 MAPK. However, the interaction of HSPA8, but not NPM1, with p-ATM or p38MPK was remarkably attenuated by apigenin, indicating that apigenin specifically hinders the interaction of HSPA8, rather than other molecules such as NPM1, with ATM and p38 MAPK (Figure 3c,d).

HSPA8, an important member of the heat shock protein 70 family, is functionally involved in chaperone-mediated autophagy, one of the main pathways of the lysosome-autophagy proteolytic system, whereas deficient protein degradation compromises cellular proteostasis and activates signaling pathways to culminate in induction of cellular senescence, a typical feature of aging ^39^. HSPA8 activates the NF-κB signaling by destabilizing IκBβ protein in the absence of lipopolysaccharide (LPS) or facilitating its nuclear translocation in the presence of LPS, an activity that can be synergized by thioredoxin domain-containing protein 5 (TXNDC5) to exacerbate the inflammatory phenotype of synovial fibroblasts in rheumatoid arthritis *via* NF-κB signaling ^40^. HSPA8 is also essential for VEGF-induced Akt phosphorylation, while downregulationn of HSPA8 abolishes Akt phosphorylation ^41^. Therefore, we speculate that HSPA8 plays a key role in the orchestrating the functional involvement of a number of factors responsible for senescence maintenance and the SASP development. To prove this hypothesis, we evaluated the expression of a series of SASP factors or senescence-associated markers. Our study indicated that in the presence of VER155008 (hereafter VER), a selective HSPA8 inhibitor, the expression of canonical SASP factors was dampened in a concentration-dependent manner (Figure 3e). Further data substantiated that cellular senescence was accompanied by IκBα degradation in the cytoplasm as well as cytoplasm-to-nucleus translocation of NF-κB subunits (p65, p50), a tendency that was generally reversed upon apigenin or VER treatment, indicating a mechanism shared by these agents in curtailing activation of the NF-κB complex (Figure 3f,g). Interestingly, treatment with VER also suppressed the occurrence of a cascade of molecular events including activation of p38 MAPK and engagement of PI3K/Akt/mTOR upon genotoxicity stress (Figure 3h).

Thus, interfering with the crosstalk between ATM and its critical targets such as HSPA8 and p38 MAPK, as exemplified by the natural agent apigenin, is able to attenuate the SASP expression, while maintaining cellular senescence or growth arrest, a distinctive feature that is basically in line with the capacity of senomorphics. As the majority of natural flavonoid polyphenols have antioxidant activities, including reducing ROS overproduction and ameliorating oxidative stress, we next asked whether apigenin has a similar efficacy in senescent cells. When exposed to a genotoxic agent (BLEO) for 8-10 d, stromal cells showed markedly elevated ROS levels when compared with their proliferating counterparts. However, in the presence of apigenin (or VER, as an experimental control), production of the ROS was significantly inhibited, largely consistent with the oxidative radical-neutralizing capacity of apigenin, although the ROS level remained largely unaffected in normal cells (Figure 3i,j).

### 2.4 Apigenin directly targets PRDX6 to attenuate its iPLA2 activity and SASP expression in senescent cells

Although the data suggested that apigenin inhibits the SASP expression through disrupting the interactions of HSPA8 with other critical factors that mediate SASP signal transduction, resulting in blocked transition of ASAP to the SASP (Figure 3c,d,e), the potential direct target(s) of apigenin in senescent cells for such a senomorphic effect remains unclear. We thus applied two affinity-based approaches, biotin-tagged apigenin (hereafter Bio-APIG) and drug affinity responsive target stability (DARTS) ^42, 43^, to reveal the potential direct target(s) of apigenin (Figure 4a and Figure S4a,b,d). In the probe-based route, upon incubation with senescent cell lysates, Bio-APIG bound to proteins before streptavidin-agarose beads were applied to pull down the Bio-APIG-protein complex for assay by MS. We observed 61 potential target proteins, which were identified with distinct enrichment by Bio-APIG but competitively inhibited by unlabeled apigenin (unique peptides ≥2, LFQ intensity ≥1.5 for enrichment or ≤0.67 for competitive inhibition), also evidenced by protein sliver staining (Figure 4b and Figure S4c). To further discover *bona fide* target proteins, we followed another approach, namely DARTS, to further substantiate the findings by target identification. The data indicated that 16 proteins were markedly upregulated upon incubation with apigenin, in relative to vehicle (unique peptides ≥2, LFQ intensity ≥1.2) (Figure 4c). Among the proteins (PRDX6, SEPT11 and YWHAE) identified by both approaches, we speculated that apigenin may directly bind to PRDX6, a hypothesis that needs to be experimentally proved (Figure 4d).

**Fig. 4.**
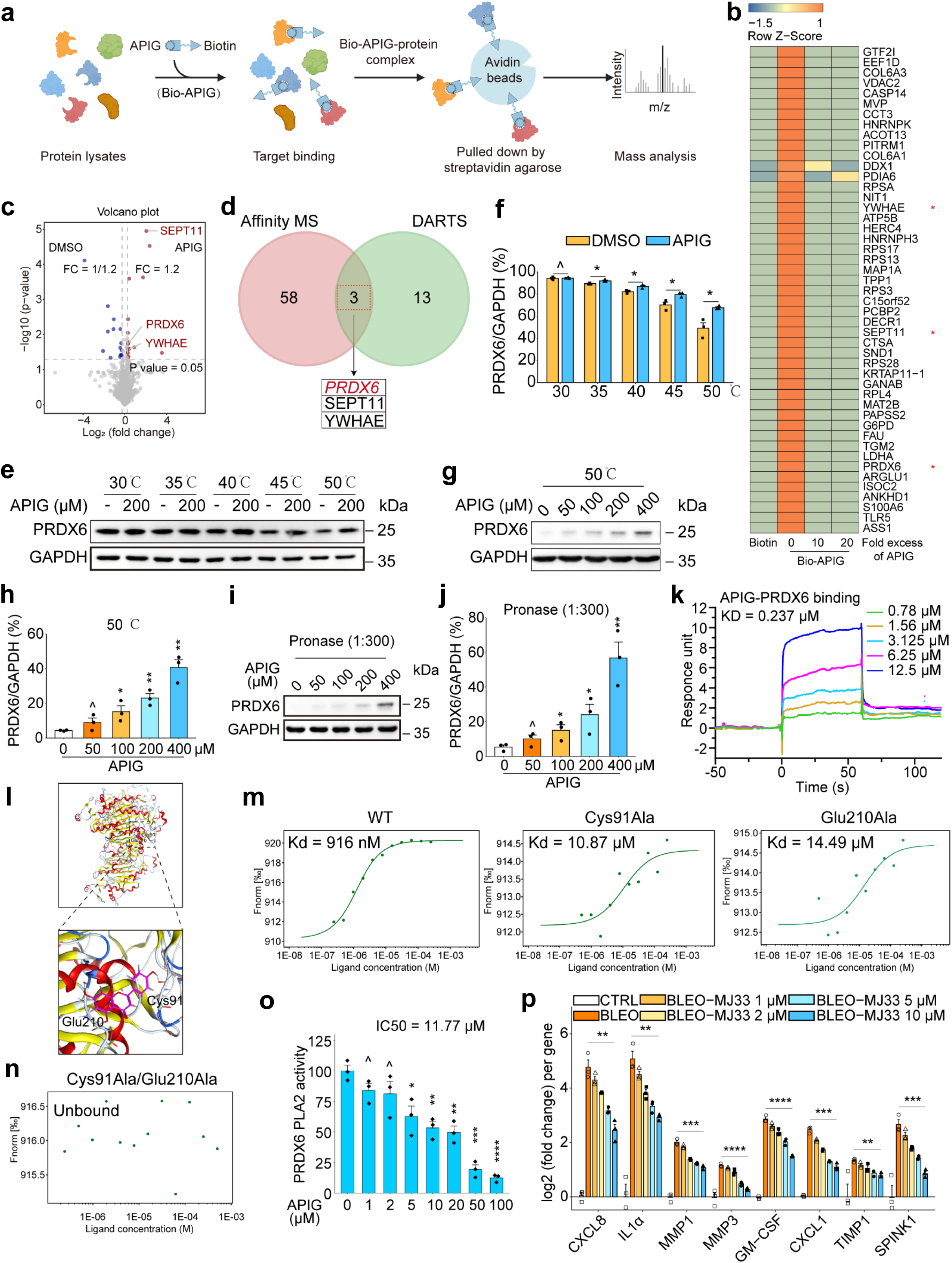
Apigenin binds to PRDX6, inhibits its iPLA2 activity and dampens the SASP development. (a) A schematic illustration of the technical procedure of affinity-based MS that applied a Bio-APIG probe for target enrichment, followed by streptavidin-agarose pull-down. This approach allows to discover target(s) of apigenin with protein lysates of senescent cells. (b) Heatmap displaying top human proteins (47) enriched by Bio-APIG probe but competitively inhibited by a 10- or 20-fold excess of unlabeled apigenin. Red stars denote proteins alternatively identified by the DARTS approach, which was technically combined with MS assay. (c) Volcano plot displaying the significantly and differentially regulated proteins (red, upregulated; blue, downregulated) in lysates of senescence cells by DARTS in the presence or absence of apigenin. Denoted proteins were also identified by Bio-APIG probing. (d) Venn diagram depicting the 3 potential target proteins identified by both Bio-APIG probing and DARTS strategies. (e) CETSA assay appraisal of the thermal stabilization of PRDX6 upon incubation with apigenin at a gradient range of temperatures from 30 to 50°C in protein lysates of PSC27 cells. (f) Comparative statistics of the thermal stabilization of PRDX6 as depicted in (e). (g) Assessment of thermal stability of PRDX6 at a gradient range of concentrations from 0 to 400 μM of apigenin when examined at a fixed temperature of 45 °C. (h) Comparative statistics of the stabilization of PRDX6 at a gradient range of concentrations of apigenin at 45°C as depicted in (g). (i) Evaluation of proteolysis stability (pronase) of PRDX6 at a gradient range of concentrations from 0 to 400 μM of apigenin when the ratio of enzyme to protein was fixed at 1:300. (j) Comparative statistics of the enzymatic stabilization of PRDX6 at a gradient range of concentrations of apigenin as depicted in (i). (k) Demonstration that PRDX6 directly binds to apigenin. In SPR assay, PRDX6 was treated by apigenin at a gradient range of concentrations from 0.78 to 12.5 μM. KD value is shown beside the traces. (l) *In silico* molecular modelling of apigenin potential bound to Cys91 and Glu210 of PRDX6. Reported X-ray diffraction microscopy structure (PDB ID: 5B6M) was applied to perform molecular docking, with Cys91 and Glu210 highlighted in structure. (m-n) The interaction between apigenin and PRDX6 or its functionally disruptive mutants (m) overexpressed in cells with GFP-tag was assessed by the cellular microscale thermophoresis analysis (MST). Alternatively, the interaction between apigenin and a double site mutant (Cys91/Ala-Glu210/Ala) of PRDX6 was analyzed (n). (o) Evaluation of iPLA2 activity upon treatment of cell lysates containing overexpressed PRDX6 with apigenin and measurement of fluorescence at a wavelength of 515 nm. (p) Quantitative assay of the expression of SASP soluble factors at transcriptional level upon senescent cell treatment with increasing concentrations of MJ33 (1-10μM), a selective PRDX6-iPLA2 inhibitor. MS, mass spectrometry. Bio-APIG, biotin-apigenin. DARTS, drug affinity responsive target stability. SPR, surface plasmon resonance. iPLA2, PRDX6 phospholipase A2. Data in f, h, j, o and p are shown as mean ± SD and representative of 3 independent biological replicates, with *P* values calculated by Student’s *t*-tests. ^, *P* > 0.05; ^, *P* > 0.05; *, *P* < 0.05; **, *P* < 0.01; ***, *P* < 0.001; ****, *P* < 0.0001.

We next interrogated the biophysical properties underpinning the interaction between PRDX6 and apigenin. As an approach to appraise protein thermal stabilization after ligand binding, cellular thermal shift assay (CETSA) was performed in a temperature range from 30 to 50 °C. We observed an increase of the stability of PRDX6 upon incubation with apigenin as compared with vehicle (Figure 4e,f). We further examined the binding effect by performing CETSA at concentrations ranging from 0 to 400 μM of apigenin either at the fixed temperature of 50°C, or the ratio of enzyme (pronase) to protein fixed at 1:300. Unsurprisingly, there appeared a remarked stability elevation of PRDX6 as reflected by both approaches in a concentration-dependent manner (Figure 4g-j). Results from surface plasmon resonance (SPR) assay, a label-free direct optical biosensor approach, showed a distinct binding of recombinant human (rh) PRDX6 in the presence of apigenin, yielding a predicted dissociation constant (KD) of 0.237 µM (Figure 4k). To further identify the residue(s) of PRDX6 responsible for the binding by apigenin, we performed *in silico* molecular modelling, with the results illustrating that apigenin establishes direct hydrogen bond connections with PRDX6 *via* the Cys91 and Glu210 of PRDX6 (Figure 4l and Figure S4e). We then performed microscale thermophoresis (MST) with exogenous PRDX6 protein linked with a GFP tag. Similar to the SPR assay, MST analysis displayed evident interaction between PRDX6 and apigenin with estimated Kd of 916 nM (Figure 4m,n). Since we speculated Cys91 and Glu210 of PRDX6 may be critical for the binding between target and ligand, we generated constructs that allow to express mutants at these 2 sites individually or simultaneously to verify our inference. In contrast to wild type PRDX6, mutation of either Cys91 or Glu210 to alanine rendered their Kd to approximately 10 times higher. However, when both sites were mutated, the binding between PRDX6 and apigenin was abrogated (Figure 4m,n and Figure S4f). Altogether, our data consistently indicated that the binding of PRDX6 and apigenin may generate a significant effect during cellular senescence.

Previous studies reported that PRDX6 has unique dual-function enzyme activities within the peroxidase family, with its phospholipase A2 (PLA2) activity rendering cells to generate arachidonic acid (AA), a molecule that is often causes inflammation ^44^. As a peroxidase, PRDX6 facilitates the conversion of H_2_O_2_ into water in the presence of NADPH, to minimize oxidation damage. Thus, we engineered a construct to overexpress PRDX6 and assessed the effect of apigenin on its capacity to remove H_2_O_2_, especially when compared with NAC, a selective inhibitor of peroxidase (Figure S4,g,h). In contrast to NAC, which remarkably reduced the peroxidase activity of purging H_2_O_2_ (IC50 = 15.5 µM), the efficacy of clearing H_2_O_2_ by apigenin was less notably compromised (IC50 = 49.5 µM) (Figure S4i), indicating that the effect of apigenin was not entirely dependent on its peroxidase activity. However, upon measurement of the PLA2 activity, we noticed a distinct reduction, implying that the capacity of apigenin to dampen the expression of pro-inflammatory factors may be attributed to its capacity to block the PLA2 activity of PRDX2 (Figure 4o). Moreover, we applied MJ33, a selective PRDX6 PLA2 inhibitor, to examine its efficacy to constrain the SASP expression, which is correlated with inflammation activity of senescent cells. We found that MJ33 treatment prominently lowered expression of the canonical SASP factors in a concentration-dependent manner (Figure 4p). Proteomics profiling showed that, unlike the vehicle, MJ33 treatment significantly inhibited the PI3K/Akt signaling pathway, further substantiating its senomorphic effect (Figure S4k). We further analyzed the protein levels of human PRDX family, which encompasses six homologs (PRDX1-6), in senescent cells in the absence or presence of apigenin treatment. We observed neither increased nor decreased expression of these homologs (Figure S4j). To further dissect the mechanism underlying the SASP downregulation by apigenin-induced suppression of the PLA2 activity of PRDX6, we performed PRDX6-mediated co-immunoprecipitation (co-IP) followed by MS analysis. The results suggested that at HSPA8 indeed does interact with PRDX6, suggesting that functional interference with either of these molecules may explain the senomorphic effect of apigenin we observed in this study (Figure S4l,m). Altogether, the data support that apigenin dampens the SASP expression through directly binging to PRDX6 and blocking its PLA2 activity.

### 2.5 Apigenin deprives cancer cells of malignancy acquired from senescent stromal cells

Many soluble SASP factors secreted by senescent stromal cells favor malignant changes of cancer cells, acting as a significant force that fuels tumor progression as demonstrated by previous studies ^26, 45–48^. Herein, we examined the capacity of the SASP factor-containing conditioned media (CM) in promoting cancer cell proliferation, a basic feature of activated stroma in tumor microenvironment (TME). The stromal cell line PSC27 was induced senescent by BLEO, a chemotherapeutic agent frequently used in cancer clinics, in the presence or absence of apigenin, with the CM collected 8-10 d afterwards and used to incubate prostate cancer (PCa) cell lines (PC3, DU145, M12, LNCaP) (Figure S5a). Not surprisingly, the proliferative potential was substantially enhanced in all PCa lines after treatment with the CM from senescent stromal cells, a tendency consistent with increased migration and invasion of these cells (Figure 5a-c). To exclude the potential effects of proliferation on observed cell migration, we performed wound healing assays in a shorter period, namely 18 h after cell scratch. Apigenin exhibited a remarkable ability to inhibit cell migration, even though no significant increase of proliferation was observed within such a timeframe. However, the malignant phenotype were markedly weakened upon apigenin treatment (Figure 5a-c).

**Fig. 5.**
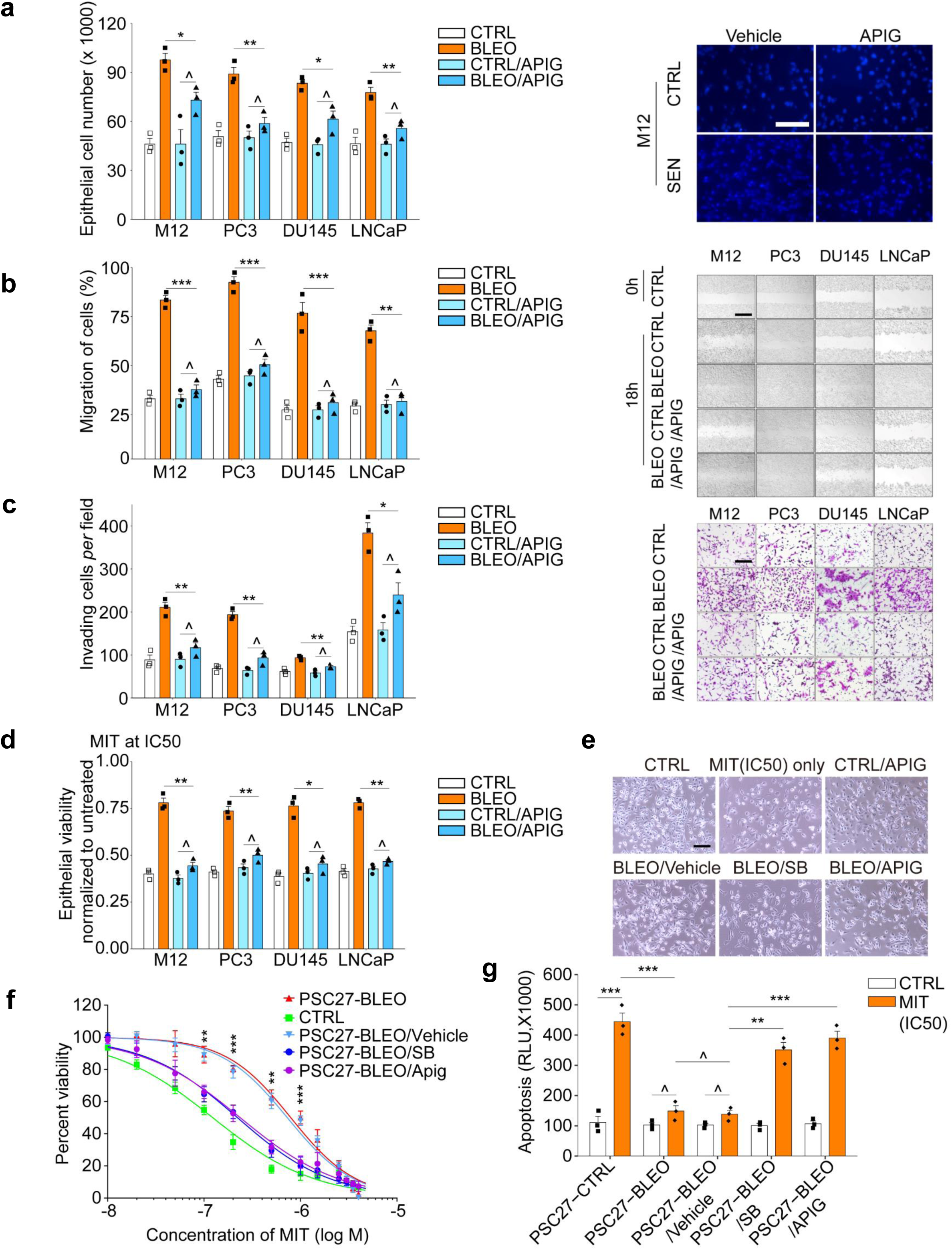
Apigenin reduces PCa cell malignancy conferred by senescent stromal cell-derived conditioned media. (a) Proliferation assay of PCa cells incubated with different types of CM for 3 days. Left, statistics. Right, representative images. Scale bar, 50 μm. (b) Migration appraisal of PCa cells incubated with several types of CM for 18 hours. Scale bar, 200 μm. (c) Invasiveness measurement of PCa cells across collagen-based transwell membrance incubated with different types of CM for 3 days. Scale bar, 200 μm. (d) Chemoresistance assay of PCa cells treated with MIT while being cultured with different types of CM. MIT was given at a pre-determined half-maximal inhibitory concentration (IC50) for each cell line. (e) Examination of cell viability in culture upon exposure to IC50 concentrations of MIT. Representative images are shown. Scale bar, 50 μm. (f) Dose-response curves (nonlinear regression/curve fit) of PC3 cells treated with different types of CM, and concurrently exposed to a wide range of concentrations of MIT. Data were plotted on an exponential scale, with cell viability assessed relative to the untreated group and calculated as a percentage. (g) Signal readings from caspase-3/7 activity assay of PC3 cells were plotted as relative luminescence units (RLUs). PCa, prostate cancer. CM, conditioned media. MIT, mitoxantrone. SB, SB203580 (a p38MAPK inhibitor that suppresses the SASP expression). Data in a, b, c, d, f and g are shown as mean ± SD and representative of 3 independent biological replicates, with *P* values calculated by Student’s *t*-tests. ^, *P* > 0.05; ^, *P* > 0.05; *, *P* < 0.05; **, *P* < 0.01; ***, *P* < 0.001; ****, *P* < 0.0001.

Our previous studies revealed that genotoxicity-damaged stromal cells in TME hold remarkable potential to confer resistance on remnant cancer cells, as evidenced by the SASP factors WNT16B, AREG and EREG ^26, 47, 49^. Whether or not the ability of the SASP factors to enhance cancer chemoresistance can be reduced by apigenin remains yet unknown. We found that the viability of cancer cells was markedly elevated upon co-culture with CM derived from senescent stromal cells, but diminished almost to the basal level as compared to their normal stromal cell counterparts upon treatment with apigenin (Figure 5d). Such a declined cancer cell viability may be explained by elevated pro-apoptotic effect of apigenin on cancer cells *via* its SASP-dampening competence, as reflected by increased activity of caspase 3/7 (Figure 5e). The capacity of aigenin in counteracting chemoresistance were also in line with the distinct shift of cell survival curves, which was based on the percentage of cell viability against MIT concentrations in the range of 0.1-10 μM, a clinically relevant window of circulating concentrations in cancer patients ^50^ (Figure 5f). To expand, we used docetaxel (DTX), another chemotherapeutic agent, to further confirm the capacity of apigenin in preventing chemoresistance. Replacement of MIT with DTX largely phenocopied the case of MIT by decreased apoptosis and increased chemoresistance, while both were considerably suppressed by apigenin (Figure S5b-d). These results suggest that senescent stromal cell-derived CM conferred cancer cells with remarkable therapeutic resistance, a tendency that was attenuated upon apigenin treatment of stromal cells.

Metformin, a widely used antidiabetic drug, is a repurposable agent and can exert multiple beneficial effects including reduction of the SASP without causing substantial cytotoxicity to senescent cells ^51, 52^, thus holding the potential as a senomorphic agent. The results from our metformin-involving assays generally phenocopied those derived from apigenin assays in lowering cancer cell malignancy by downregulating the activity of stromal cell CM, indicating a common pattern shared by both agents (Figure S5e-h). Thus, pharmacologically targeting the SASP development with a senomorphic agent such as apigenin, can substantially deprive cancer cells of these stromal CM-conferred gain-of-functions, specifically resistance to a chemotherapeutic drug. These findings imply the possibility of developing relevant strategies to improve the efficacy of current anticancer regimens.

### 2.6 Apigenin combined with chemotherapy effectively reduces chemoresistance

Given the efficacy of apigenin in attenuating the SASP expression in senescent stromal cells and restraining malignant phenotypes of cancer cells *in vitro*, it is tempting to determine whether this agent holds a potential to control senescence-related pathologies *in vivo*. To this end, PSC27 sublines were mixed together with PC3, a typically malignant prostate cancer line, at a pre-optimized ratio (1:4) to generate tissue recombinants prior to subcutaneous inoculation to the hind flank of mice with non-obese diabetes and severe combined immunodeficiency (NOD– SCID). The tumor sizes were determined 8 weeks later for pathological appraisal. In comparison to tumors made up of PC3 and PSC27^Naive^, xenografts comprising PC3 and PSC27^SEN^ were remarkably enlarged. However, treatment of PSC27^SEN^ cells with apigenin prior to tissue recombinant construction substantially reduced tumor volumes (*P* < 0.01). Notably, the efficacy of apigenin was largely reproduced by rapamycin, which exerted a durable *in vivo* effect even when senescent human cells were treated only once by such a SASP-inhibiting agent *in vitro* (Figure S6a)^53^.

To closely simulate clinical settings, we designed a preclinical regimen involving genotoxic agents and/or apigenin (Figure 6a and Figure S6b). Following a two-week period after subcutaneous implantation with generally observable tissue recombinant uptake in host animals, a single dose of placebo, MIT or apigenin was administered at the beginning of the 3^rd^, 5^th^ and 7^th^ week until the end of a 8-week regimen (Figure S6b). Although no significant benefits were observed in the apigenin group, MIT administration caused remarkable tumor shrinkage (57.8% reduction in volume), confirming the effectiveness of MIT as a chemotherapeutic agent. When apigenin was co-administered with MIT, we noticed an additional reduction in tumor size by 51.1%, corresponding to a total decrease by 74.9% as compared with the placebo group. We next queried whether cellular senescence occurred in xenografts. As expected, in the TME of PC3/PSC27 recombinant tumors, stromal cells exhibited a markedly elevated expression of canonical SASP factors including IL-1α/IL-6, CXCL8, MMP1/MMP3, ANGPTL4 and AREG, a pattern arising in parallel with the upregulation of typical senescence markers p16^INK4a^ and p21^CIP1^ in the MIT group, indicating senescence induction *in vivo*and the SASP expression upon chemotherapeutic treatment (Figure 6c,d and Figure S6c). Moreover, histologic staining indicated enhanced SA-β-Gal positivity in xenografts of mice exposed to MIT, a feature in sharp contrast to apigenin, which neither promoted nor suppressed cellular senescence (Figure 6e), a tendency in line with our results acquired from *in vitro* assays (Figure 2a). The alterations were predominantly observed in stromal cells instead of adjacent cancer cells, indicating the potential of residual cancer cell repopulation and resistance acquisition in the treatment-damaged TME. However, upon delivery of apigenin, the SASP expression was basically dampened, as evidenced by data from RNA-seq assays (Figure S6d).

**Fig. 6.**
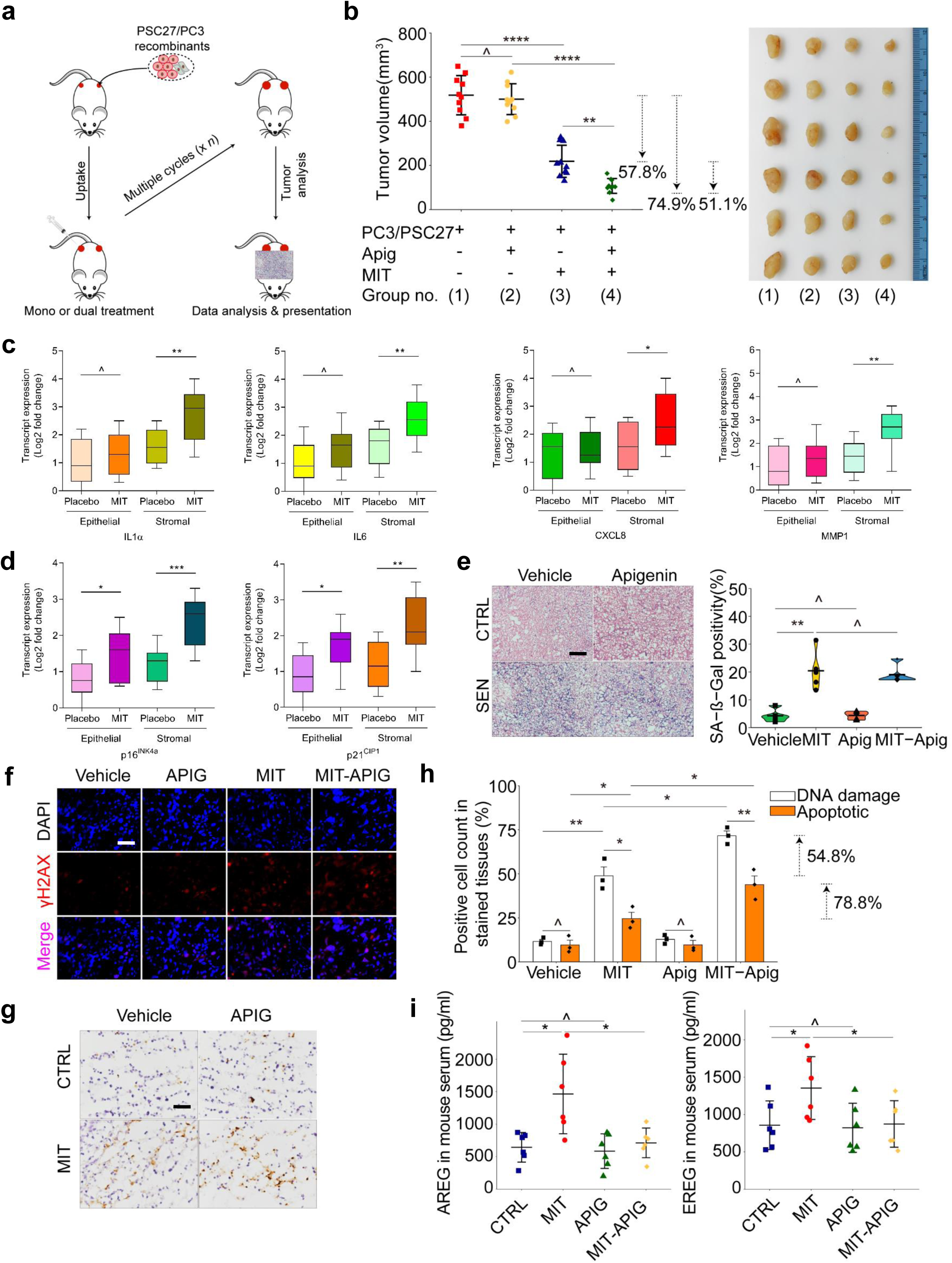
Combination of apigenin and chemotherapy improves preclinical outcomes. (a) A schematic workflow illustrating the procedure of a preclinical regimen. Two weeks after subcutaneous inoculation and *in vivo* uptake of PC3/PSC27 recombinants, severe combined immunodeficient (SCID) mice underwent either single agent or combinatorial treatment in a metronomic schedule composed of several cycles. **(b)** Statistical analysis of tumor end volumes. PC3 cells were inoculated either alone or combined with PSC27 before implanted subcutaneously to the hind flank of NOD/SCID mice, which were administered with MIT and apigenin, alone or concurrently. Left, comparative statistics. Right, representative tumor images. (c) Transcriptional analysis of several typical SASP factors in stromal cells isolated from PC3/PSC27 tumor foci, which were subjected to laser capture microdissection (LCM) isolation for stromal and cancer cells, respectively. Signals were normalized to the lowest sample in placebo group. (d) Transcriptional analysis of two canonical senescence biomarkers *p16^INK4a^* and *p21^CIP1^*. (e) Appraisal of cellular senescence in xenograft tissue by SA-β-Gal staining. Left, representative images. Right, comparative statistics. Scale bar, 100 μm. (f) Quantification of DDR signals by immunofluorescence (IF) staining for γH2AX in xenograft tissues (red, γH2AX; blue, DAPI). Scale bar, 20 μm. (g) Assessment of cell apoptosis in xenograft tissues by immunohistochemistry (IHC) staining for caspase cleaved 3 (CCL3) at the completion of treatment regimens. Biopsy samples from animals receiving a placebo served as negative controls for mice receiving treatments by MIT and/or apigenin. Scale bar, 20 μm. (h) Statistical evaluation of DNA damage and cell apoptosis in xenograft tissues. Values are shown as the percentage of cells positively stained by IF or IHC specific to γH2AX or CCL3, respectively. (i) Evaluation of the circulating levels of two typical SASP factors (AREG, left; EREG, right) in the serum of animals receiving MIT and/or apigenin treatments. DDF, DNA damage foci. Data in b, c, d, e, h and i are shown as mean ± SD and representative of 3 independent biological replicates, with *P* values calculated by Student’s *t*-tests. ^, *P* > 0.05; ^, *P* > 0.05; *, *P* < 0.05; **, *P* < 0.01; ***, *P* < 0.001; ****, *P* < 0.0001.

We next interrogated how expression of the SASP induces the therapeutic resistance of cancer cells. To explore the underlying mechanism, we dissected tissues from animals 7 days after treatment initiation, a timepoint just prior to the emergence of resistant colonies. Contrasting the placebo group, MIT treatment *per se* enhanced DNA damage response (DDR) and caused cell death particularly apoptosis in cancer cells, while the administration of apigenin alone triggered neither alterations, implying limited effectiveness of apigenin when used as a single agent for tumor treatment (Figure 6f-h). Upon co-administration of apigenin and MIT, intensities of both DNA damage and apoptosis were further elevated, indicating increased cytotoxicity upon combinatorial therapy. We further observed enhanced caspase 3 cleavage (CCL), a typical cellular apoptosis marker, when apigenin was combined with MIT (Figure 6g). Upon ELISA analysis of circulating AREG and EREG, typical SASP factors, we noticed that MIT treatment resulted in a remarkably elevated circulating levels of these factors, a tendency that was essentially reversed upon apigenin administration (Figure 6i).

To establish the safety and effectiveness of our therapeutic strategy, a systematic appraisal of the physiology of experimental animals was performed. Of note, gross assessment data indicated that both single and co-administration were well tolerated, as evidenced by the absence of body weight loss between different groups during the entire regimens (Figure 6e). Further, no remarkable fluctuation was observed in the serum level of creatinine, urea and metabolic status including alkaline phosphatase (ALP) and alanine aminotransferase (ALT) activities, which represent a set of biochemical measurements of major organs such as liver and kidney (Figure 6f). Hence, co-administration of a senomorphic agent and canonical chemotherapy holds the potential to achieve maximal anticancer effects while minimizing severe systemic toxicities.

### 2.7 Preclinical treatment by apigenin alleviates age-related physical dysfunction and cognitive decline

In the tissue microenvironment, the SASP is a major executor of the paracrine effects of senescent cells and mediates various local and systemic biological effects, including those underlying pathologies and impacting overall healthspan ^54^. To further determine the efficacy of apigenin in regulating senescence-related *in vivo* changes including organismal aging, we chose to employ irradiation-challenged mice for aging-related studies. Briefly, wild type (WT) mice were exposed to a sublethal dose of whole body irradiation (WBI) to induce substantial senescence within tissues, followed by senotherapeutic intervention with apigenin or vehicle, the latter a therapeutic control. As a result, animals experiencing WBI manifested an aberrant physiological profile, as exemplified by apparently graying fur (Figure 7b). However, this condition was markedly improved upon treatment by apigenin.

**Fig. 7.**
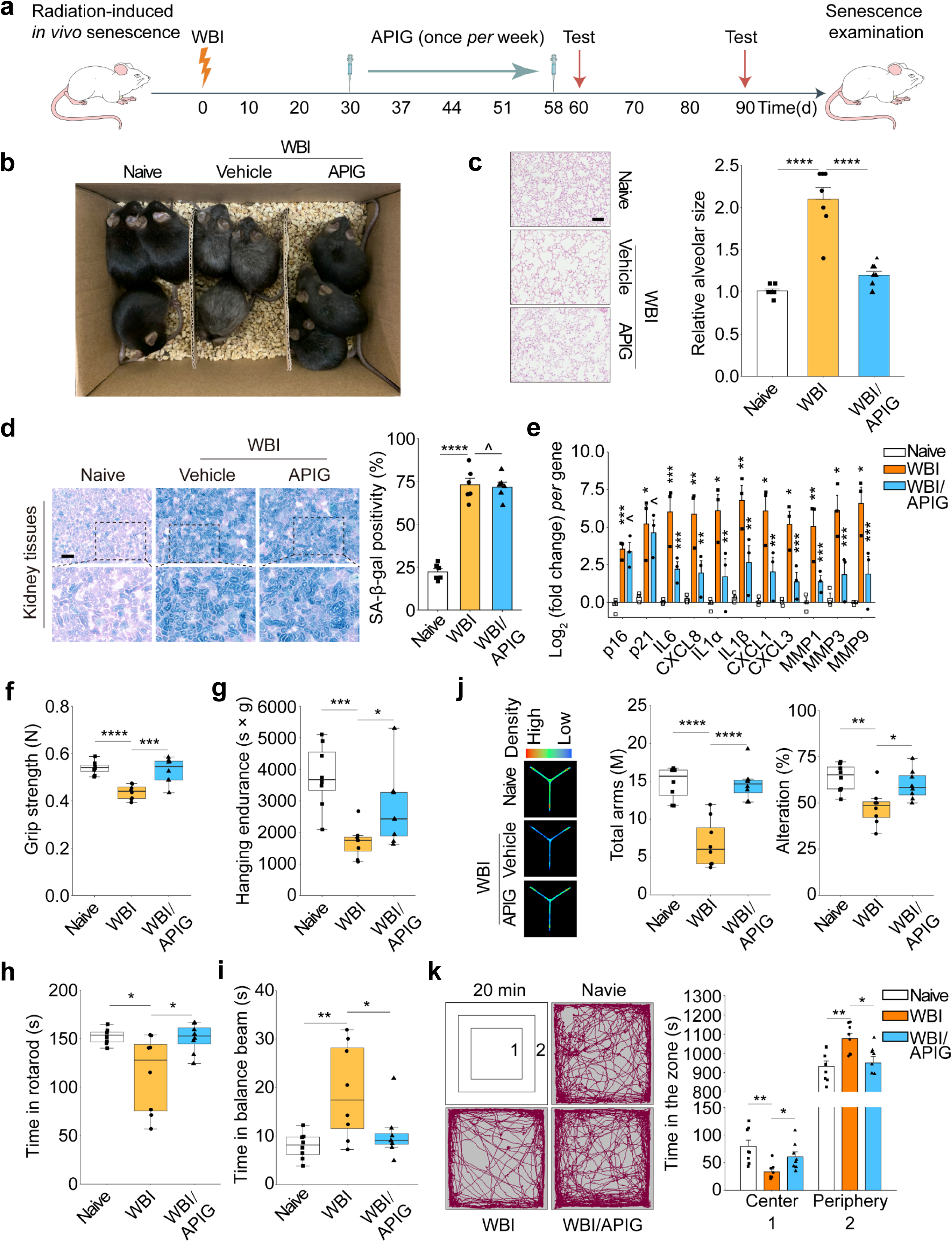
Apigenin administration improves physical function of mice prematurely aged after WBI treatment. (a) Schematic illustration of a preclinical procedure for C57BL/6J animals undergoing WBI treatment and subject to physical function tests. (b) A representative snapshot of the whole body profile of experimental animals. Naïve mice, WBI mice experiencing vehicle treatment and WBI mice receiving apigenin administration, respectively, were recruited to the study. (c) H&E staining to dissect the morphology of lung tissues from mice as depicted in (b). Scale bar, 100 μm. (d) Representative images of SA-β-Gal staining (left) and quantification (right) from kidney tissues from each group. Scale bar, 5 μm. (e) Quantitative assessment of the transcript expression of senescence markers (p16^INK4a^ and p21^CIP1^) and several typical SASP factors in kidney tissues. Signals were normalized to samples of the naïve group *per* gene. (f-i) Quantitative appraisal of physical functions, including grip strength (f), hanging endurance (g), time in rotarod (h) and time needed for crossing the balance beam (i), all typical physical capacities of animals depicted in (a). n = 8. (j) Y maze test of mice. n = 8. (k) Evaluation of the short-term memory of animals as depicted in (a) *via* open field test during a 20 min exploration. ‘1’ and ‘2’ denote the ‘central’ and ‘peripheral’ zones, respectively, of the open field (top left). The representative trajectory (left) and the quantification analysis (right) indicate the time animals spent in two zones. n = 8. Each data point represents an individual mouse. n represents the number of animals *per* group. WBI, whole body irradiation. H&E, hematoxylin and eosin. Data in c-k are shown as mean ± SD and representative of 3 independent biological replicates. *P* values were calculated by Student’s *t* tests. ^, P > 0.05; ^, *P* > 0.05; *, *P* < 0.05; **, *P* < 0.01; ***, *P* < 0.001; ****, *P* < 0.0001.

**Fig. 8.**
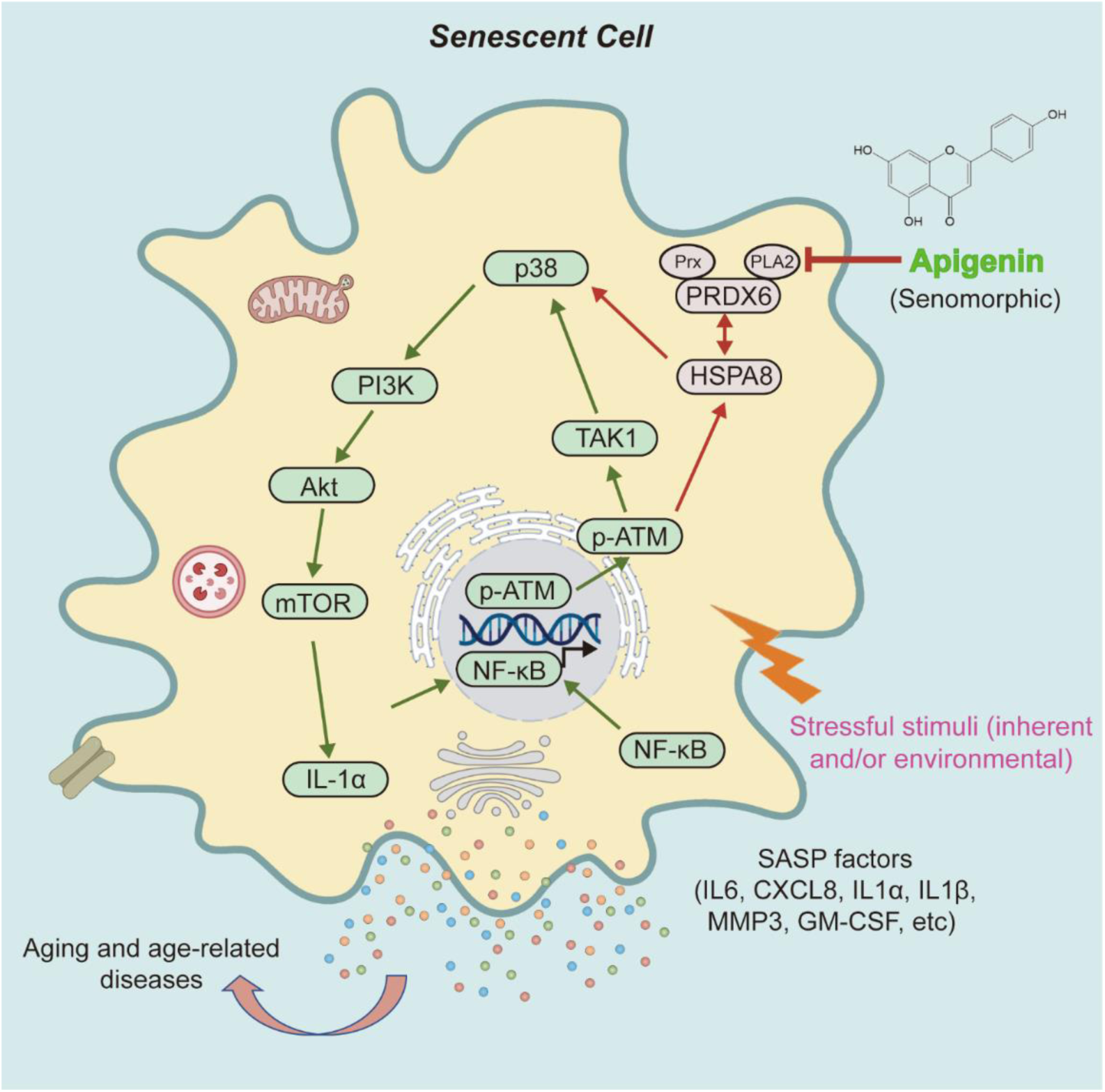
A working model of apigenin as a novel senomorphic agent to target senescent cells and exploration of its potential in future antiaging pipelines. Senescent cells actively synthesize and secrete a myriad of SASP factors, which encompass multiple soluble proteins including pro-inflammatory interleukins, chemokines and growth factors. The hypersecretory feature of senescent cells, namely the SASP, affect the physiology of neighboring cells and causing chronic inflammation in the tissue microenvironment. As a naturally derived small molecule compound, apigenin directly targets PRDX6 to restrain its PLA2 activity, disrupting HSPA8 activation and interfering with its interactions with ATM and p38 MAPK in senescent cells. Through restraining the SASP expression, apigenin effectively minimizes the impact of senescent cells on tissue homeostasis, preventing organ function degeneration and ameliorating multiple age-related conditions.

Age-dependent expansion of pulmonary alveolus, an important tissue structural alteration that contributes to pulmonary dysfunction, was found in WBI-treated mice, but much lightened in animals receiving apigenin treatment (Figure 7c). Our data suggest a benefit of apigenin supplementation in preventing the appearance of age-related pulmonary abnormality, as changes in the composition of the airways and the alveoli may cause respiratory dysfunction and eventually promote chronic lung disorders during chronological aging ^55^. Upon histological assessment of spleen tissues of aged mice, we observed atrophy and disarrangement of white pulp, the major immune organ comprising various immune cell types, including B cells, DC cells, CD4^+^ and CD8^+^ T cells, abnormal changes of which may cause age-associated immune dysfunction (Figure S7a). However, apigenin exhibited a prominent capacity to improve the structural profile of spleen tissues by preventing such abnormalities.

Subsequent analysis indicated the emergence of senescent cells in several major organs, as demonstrated by increased SA-β-Gal positivity in liver and kidney tissues of these animals post WBI-treatment (Figure 7d and Figure S7a,b). As compared with vehicle-treated mice, the tendency of SA-β-Gal staining positivity remained largely unchanged in the apigenin group, suggesting that treatment with such a senomorphic agent cannot prevent cellular senescence in these animals. We further noticed that apigenin administration failed to downregulate the expression of key senescence markers including p16^INK4a^ and p21^CIP1^ (Figure 7e). The data was further validated by immunofluorescence analysis of p21 expression in the tissue microenvironment (Figure S7c). However, in contrast, a subset of typical SASP factors including IL6, CXCL8, IL1α, IL1β, CXCL1/3 and MMP1/3/9, which were significantly upregulated after exposure of mice to WBI, exhibited remarkable decline in animals treated by apigenin (Figure 7e). This observation was largely consistent with the data acquired from mice involved in chemotherapeutic regimens (Figure S6d). Alternatively, we examined the biospecimens at protein level. Our data indicated that the upregulation of the canonical SASP marker IL6 in tissues such as those within the kidney after animals underwent WBI treatment, was remarkably reversed in animals treated by apigenin (Figure 7d,e). Upon performance of ELISA assays to measure the circulating levels of AREG and EREG, we observed an apparent elevation of circulating concentration of these typical SASP factors in irradiation-challenged animals, a tendency that was largely counteracted upon apigenin administration (Figure S7f,g).

Next, we assessed whether preclinical interventions altered the physical functions of these animals. Unsurprisingly, mice experiencing WBI manifested reduced exercise performance and muscle strength, as gauged by grip strength, hanging endurance, rotard rod duration and balance cross tests (Figure 7f-i). However, decline in each of these activities were partially but significantly reversed upon treatment of prematurally aged animals with apigenin, as compared with the vehicle group. We further queued whether apigenin could generally improve health conditions including short term memory and anxiety extent of animals. Notably, we observed a basic recovery of short-term memory in aged animals that received apigenin treatment, as evidenced by data from Y-maze tests (Figure 7j). Data from open field assays indicated that apigenin prolonged the duration of exploring the central zone of a wide arena, suggesting that the anxiety of aged mice was remarkably alleviated in aged mice as compared with the vehicle group (Figure 7k). The data suggest that apigenin administration represents an effective approach to alleviate physical dysfunction and avoid cognitive impairment during premature aging, thus promoting overall health conditions.

To establish the safety of our interventional strategy in these immunocompetent animals (C57BL/6J), a systemic and routine evaluation of blood cell counts and serum biochemical indices was performed. As a result, we observed a slight decrease in the counts of both white blood cells (WBCs) and red blood cells (RBCs) in WBI-treated mice (Figure S7h-j). Although the difference appeared insignificant, it reflected a partial decline of hematopoietic capacity and immune competency. Upon apigenin administration, these indices remained generally unchanged, validating the safety of apigenin treatment in the hematopoiesis system (Figure S7h-l). As a further evidence, we found no obviously abnormality in the serum levels of creatinine, urea, and activities of ALP and ALT, typical liver and renal biochemical parameters (Figure S7m-p). Altogether, our preclinical data support that the therapeutic regimen involving a senomorphic agent, such as apigenin, in prematurely aged mice holds a prominent anti-aging potential without causing systemic cytotoxicity.

## DISCUSSION

Aging affects the functional integrity of various organ types and leads to chronic degenerative pathologies ^2, 56^.The expanding human aging population in most developed countries has generated significant societal and economic costs, making development of antiaging therapeutics an outstanding need. One promising avenue is to explore the potential of natural products, particularly botanical extracts, which are rich in phytochemical constituents with a capacity to serve as health-promoting agents ^57^. Increasing lines of evidence suggest that naturally derived compounds targeting cellular senescence hold the potential to improve healthspan and extend lifespan of various mammalian species, specifically rodent and human ^11, 14, 15, 58^. Cellular senescence is a state representing a permanent cell cycle arrest while cells remain viable, but it contributes to organ degeneration and age-related diseases when the threshold senescent cell accumulation is exceeded in the time course of aging ^8, 59^.

The pathogenic roles of senescent cells have been well recognized, with more recent scientific and medical efforts aimed at making them a potentially treatable target ^16, 60, 61^. As a critical mediator of senescent cell-associated pathophysiological effects, the SASP can be effectively targeted through two principal strategies, radical clearance of senescent cells with senolytics and specific inhibition of the SASP *per se* using senomorphics, together termed as senotherapeutics ^59, 62^. Promising results have been observed in preclinical studies, and more importantly, in clinical trials, which involve administration of senolytics for age-related disorders including diabetic kidney disease (DKD), idiopathic pulmonary fibrosis (IPF), sight-threatening diabetic macular edema (DME) and early stage Alzheimer’s disease (AD) ^12, 13, 15, 63^. In contrast, a major advantage of senomorphics, for example, rutin and resveratrol, is the ability to selectively restrain the SASP expression, but without the need to eliminate senescent cell populations *in vivo* ^29, 32^. This approach, although considered pharmacologically milder than senolytics, can effectively avert the potentially harmful effects of the SASPs, while allowing senescent cells to stay in the tissue microenvironment, which may play essential roles such as impeding tumor development by enhancing immunosurveillance and other physiologically essential activities ^64^.

In this study, we performed *in vitro* screening of a NMA library to search for candidates of natural senotherapeutics and noticed a small molecule compound apigenin, which exhibits a prominent senomorphic effect. Treatment of senescent cells with apigenin can abrogate the expression of a wide spectrum SASP, a key player that mediates the occurrence and progression of most, if not all, age-related pathologies ^4^. Data from proteome-wide profiling suggested the possibility of HSPA8 to interact with ATM and p38 MAPK, two molecules known to be implicated in the transduction of intracellular signaling to allow the SASP development upon cellular senescence ^65, 66^. Further studies indicated that apigenin interferes with the crosstalk between ATM and HSPA8, as well as that between HSPA8 and p38 MAPK, thus blocking more than one signaling sections that functionally mediate the SASP progression, which usually occurs in a cascade manner in senescent cells. Therefore, apigenin abrogates the transition of the ASAP, an acute response of cells after exposure to stress, towards a full scale SASP, a chronic and pro-inflammatory phenotype. Indeed, such a functional efficacy is reminiscent of 5Z-7-oxozeaenol, a TAK1 inhibitor, which diminishes the SASP expression by restraining the activity of TAK1, a kinase that is involved in the ATM-TRAF6-TAK1 axis during acute DNA damage response and subsequently orchestrates the engagement of p38 MAPK and PI3K/Akt/mTOR pathways to support a persistent SASP signaling ^46^.

PRDX6 is a unique 1-Cys member of the peroxiredoxin family, which comprises six highly conserved antioxidant enzymes (PRDX1-PRDX6) featuring a cysteine (cys) residue involved in peroxide reduction ^67^. PRDX6 has peroxidase, acidic calcium-independent phospholipase A2 (aiPLA2) and lysophosphatidylcholine acyltransferase (LPCAT) activities, and is important for the maintenance of lipid peroxidation repair, inflammatory signaling and antioxidant damage response ^68^. Although PRDX6 is extensively investigated in brain disorders such as AD and Parkinson’s disease (PD), the functional role of PRDX6 in cellular senescence and its associated phenotypes remains largely unknown. In this study, we first evaluated the effect of apigenin on cellular capacity of scavenging H_2_O_2_. The data suggested that apigenin reduced the ability of cells to clear H_2_O_2_, but in a less effective manner than that of NAC, a selective inhibitor of peroxidase activity, suggesting that the efficacy of apigenin may not rely on its peroxidase activity. Upon alternative dissection, which was conducted on the PLA2 activity, we revealed that the capacity of apigenin to dampen pre-inflammatory factor expression was indeed correlated with suppressing the PLA2 activity of PRDX6. The finding was further substantiated by assays involving MJ33, a selective PRDX6 PLA2 inhibitor, which remarkably decreased the SASP expression in a concentration-dependent manner. Through PRDX6-mediated Co-IP assay followed by MS profiling, we uncovered the interaction of HSPA8 with PRDX6, two molecules functionally involved in mediating the senomorphic effect of apigenin. Mechanistically, targeting senescent cells by apigenin interferes with PRDX6 PLA2 activity, affecting HSPA8 activation and abrogating downstream events including its interactions with both ATM and p38 MAPK.

Given the prominent potential of apigenin in dampening the SASP expression, we examined the capacity of this compound in restraining the malignancy of cancer cells. Experimental results indicated that almost all the gained functions, namely a series of enhanced malignant phenotypes of PCa cells (a cancer cell model used in this study), were markedly reduced by apigenin. Importantly, augmented chemoresistance of cancer cells conferred by their senescent cell counterparts, was largely deprived in the presence of apigenin, implying the possibility of applying apigenin in combinatorial anticancer treatments. We further extended these findings to a mouse model. The data showed that biweekly treatment with apigenin *via* i.p. significantly inhibited the SASP expression *in vivo* of mice carrying PCa xenografts, resulting in significantly decreased tumor volumes at the end of therapeutic regimens. More importantly, preclinical administration of apigenin to mice prematurely aged by WBI remarkably ameliorated age-related symptoms. Typical geriatric symptoms, including fur modification, alveolar volume expansion, muscle strength loss, hanging endurance reduction and beam balance crossing difficulty, were substantially improved in aged animals receiving administration of apigenin. Interestingly, cognitive impairment, a neurodegenerative deficit observed in these mice were also markedly prevented by apigenin. Although further studies remain necessary to establish the possible benefits of apigenin in improving other age-related health conditions, our preclinical evidence suggests a prominent role of apigenin in mitigating physical dysfunctions of aged animals, Principally by targeting the senescence-associated inflammatory phenotype, namely the SASP.

One of the strengths of this study is that we determined the role of apigenin against cellular senescence occurring in cell lines representative of different organ origins as well as upon different forms of senescence induced by alternative stimuli, including the cases of RS, OIS and TIS. In this study, we mainly focused on the case of TIS, as senescence accumulation within otherwise healthy tissues is an off-target effect of most chemotherapies ^46, 69^. We used BLEO, a genotoxic agent frequently applied in cancer clinics, for *in vitro* senescence induction and phenotype assessments. Alternatively, we employed MIT, another chemotherapeutic agent that induces senescence by acting as an inhibitor of topoisomerase II to cause DNA damage ^70^. Administration of MIT to experimental mice triggered significant DDR and cellular senescence *in vivo*. However, apigenin further promoted both DNA damage and cell apoptosis in tumor foci, suggesting that cancer chemoresistance conferred by *in vivo* senescence can be remarkably weakened by apigenin. To date, most senotherapeutic or anti-senescence studies chose to use senolytics in design and complementation of preclinical interventions against age-related pathologies, as removal of senescent cells does result in rejuvenated tissues and organs, a benefit that has been demonstrated by multiple studies including those apply natural agents ^11, 71^. Although it is well established that of senescent cell accumulation leads to persistent release of SASP and contributes to low-grade inflammation fueling age-related degenerative diseases ^72^, much less efforts are invested to explore the therapeutic potential of senomorphics in long term age-related settings.

Apigenin is a flavonoid (4’,5,7-trihydroxyflavone) and holds substantial promise as a preventative agent against chronic disorders. Holding a radical scavenging capacities, apigenin displays neuroprotective, anti-inflammatory and antioxidant efficacies ^73^. In contrast, its therapeutic potential in targeting senescent cells and minimizing senescence-related pathological impacts *via* blockage of the SASP expression, remains largely overlooked. In this study, we unmasked a new mechanism by which apigenin can specifically alter the senescence phenotype, specifically curtailing the SASP expression. We further demonstrated the benefits of using apigenin to intervene age-related pathologies, thus opening a new avenue to exploit such a small molecule compound as a novel senomorphic agent. Taken together, our study supports that apigenin, a plant-derived flavonoid, can effectively ameliorate age-related disorders, as exemplified by increased anticancer efficacy, improved physical dysfunction and enhanced cognitive behavior of prematurely aged mice. We present a series of proof-of-concept evidence to show that a senomorphic agent can exert significant geroprotective effects with a relatively low dose and a favorable safety profile. This study provides a pilot and efficient platform for future research and development of natural compounds as senotherapeutics to prevent, delay and ameliorate a number of age-related pathologies.

## METHODS

### Cell culture

The primary normal human prostate stromal cell line PSC27 was a courteous gift from Dr. Peter Nelson (Fred Hutchinson Cancer Center) and maintained in stromal complete medium as described previously ^26^. Human fetal lung stromal lines WI38 and IMR90 were from ATCC and cultured with F-12K medium supplemented with 10% FBS. Prostate cancer epithelial cell lines PC3, DU145 and LNCaP, and human embryonic kidney line 293T were from ATCC and routinely cultured with RPMI 1640 (10% FBS). Prostate cancer epithelial line M12 was kindly provided by Dr. Stephen Plymate (University of Washington), which originally derived from the benign line BPH1 but phenotypically neoplastic and metastatic ^74^. All lines were routinely tested for mycoplasma contamination and authenticated with STR assays.

### Cell treatments

Stromal cells were grown until 80% confluent (CTRL) and treated with 50 μg/mL bleomycin (BLEO). Cells were rinsed briefly with PBS and allowed to stay for 8-10 days prior to performance of various examinations. To perform drug screening for potential senolytics, natural candidates (totally 66 in the NMA library) (Selleckchem L8300-TCM subset) were tested each at 3 μg/mL on the survival of 5.0 × 10^3^ control and senescent cells for 3 d. Subsequent evaluation of the effects of these candidate agents (each applied at 1 μg/mL) was conducted to test the extent of SASP inhibition (assay for senomorphics). For those with a remarkable potential to act as effective and safe senomorphics, further assessments were followed. The phytochemical agent apigenin was examined in the range of 1 to 10 μM, with 10 μM determined to be a minimally optimal concentration for senomorphics activity. The positive control senolytics ABT-263 and PCC1 were applied at 1.25 μM and 50 μM, respectively. The small molecule inhibitor SB203580 of p38MAPK was used at 10 μM, while the PRDX6-PLA2 inhibitor MJ33 was employed at 10 μM, to treat senescent cells in culture before collection for lysis and expression analysis.

### HPLC-QTOF-MS/MS analysis

The standard HPLC-QTOF-MS/MS (high performance liquid chromatograph conjugated with quadrupole time-of-flight-tandem mass spectrometer) analysis was performed on Nexera X2 LC-40 (SHIMADZU) coupled to AB SCIEX TripleTOF 6600 LC/MS/MS system (SCIEX), data processed with Analyst TF (v 1.7.1). Briefly, the chromatographic separation was carried out on an Accucore C30 column (2.6 um, 250 × 2.1 mm, Thermo) at room temperature. The mobile phase solvent comprises 1% acetic acid in water (solvent A) and 100% methanol (solvent B). The multistep linear gradient solvent system started with 5% B and increased to 15% B (5 min), 35% B (30 min), 70% B (45 min), 70% B (49 min) and 100% B (50 min), held at 100% B for 10 min, and decreasing to 5% B (60 min). At the initial and last gradient step, the column was equilibrated and maintained or washed for 10 min with 5% B. The flow rate was 0.2 ml/min, while the injection volume of samples was 10 μl. Detection was performed using a TripleTOF 6600 qTOF mass spectrometer (AB SCIEX) equipped with an ESI interface in negative ion mode. Operation conditions of the ESI source were as follows: capillary voltage, -4000 V; drying gas, 60 (arbitrary units); nebulization gas pressure, 60 psi; capillary temperature, 650°C; collision energy, 30. The mass spectra were scanned from 50 to 1200 m/z, at an acquisition rate 3 spectra *per* second. Data acquisition and analyses were performed with Analyst TF (AB SCIEX, v 1.7.1) software.

### Bulk RNA-seq and bioinformatics

Total RNA samples were prepared from senescent PSC27 cells cultured with regular DMEM or DMEM containing apigenin (10 μM) for 3 consecutive days. Sample quality was validated by Bioanalyzer 2100 (Agilent), and RNA was subjected to sequencing by Illumina NovaSeq 6000 with gene expression levels quantified by the software package RSEM (https://deweylab.github.io/RSEM/). Briefly, rRNAs in the RNA samples were eliminated using the RiboMinus Eukaryote kit (Qiagen, Valencia, CA, USA), and strand-specific RNA-seq libraries were generated using the TruSeq Stranded Total RNA preparation kits (Illumina, San Diego, CA, USA) according to the manufacturer’s instructions before deep sequencing.

Pair-end transcriptomic reads were mapped to the reference genome (GRCh38.p14) (Genome Reference Consortium Human Build 38; INSDC Assembly <u>GCA_000001405.28</u>, 12.2013) (http://asia.ensembl.org/Homo_sapiens/Info/Index) (ensembl_105) with reference annotation from GENCODE v42 using the Bowtie tool. Duplicate reads were identified using the picard tools (1.98) script mark duplicates (https://github.com/broadinstitute/picard) and only non-duplicate reads were retained. Reference splice junctions are provided by a reference transcriptome (Ensembl build 73) ^75^. FPKM values were calculated using Cufflinks, with differential gene expression called by the Cuffdiff maximum-likelihood estimate function ^76^. Genes of significantly changed expression were defined by a false discovery rate (FDR)-corrected *P* value < 0.05. Only ensembl genes 73 of status “known” and biotype “coding” were used for downstream analysis.

Reads were trimmed using Trim Galore (v0.6.1) (http://www.bioinformatics.babraham.ac.uk/projects/trim_galore/) and quality assessed using FastQC (v0.10.0) (http://www.bioinformatics.bbsrc.ac.uk/projects/fastqc/). Differentially expressed genes were subsequently analyzed for enrichment of biological themes using the DAVID bioinformatics platform (https://david.ncifcrf.gov/) and the Ingenuity Pathways Analysis program (http://www.ingenuity.com/index.html). Raw data of bulky RNA-seq were deposited in the NCBI Gene Expression Omnibus (GEO) database under the accession code <u>GSE273159</u>.

#### Venn diagrams

Venn diagrams and associated empirical *P*-values were generated using the USeq (v7.1.2) tool IntersectLists ^77^. The *t*-value used was 22,008, as the total number of genes of status “known” and biotype “coding” in ensembl genes 73. The number of iterations used was 1,000.

#### RNA-seq heatmaps

For each gene, the FPKM value was calculated based on aligned reads, using Cufflinks ^76^. Z-scores were generated from FPKMs. Hierarchical clustering was performed using the R package heatmap.2 and the distfun = “pearson” and hclustfun = “average”.

### Mapping of unique interactors with high-throughput datasets

Profiling of ATM- and p38 MAPK-interactive molecules was performed with BioGRID (v4.4.237), a biomedical interaction repository with data compiled through comprehensive curation efforts and used as a public database archiving and disseminating genetic and protein interaction data from all major model organisms and humans (thebiogrid.org) ^78, 79^. BioGRID searches 85,467 publications for 2,800,228 protein and genetic interactions, 31,144 chemical interactions and 1,128,339 post translational modifications from major model organism species, with human selected as the target species throughout this study.

### Cellular thermal shift assay (CETSA)

To assess whether apigenin targets PRDX6 protein, CETSA-immunoblot experiments were carried out. CETSA is used to examine the binding efficiency of drug and protein. Briefly, PSC27 cells in culture were trypsinized and precipitated by centrifugation, lysed on ice with RIPA buffer containing protease inhibitor cocktail and centrifuged at 14,000 × g at 4 °C for 20 min. Each lysate was then equally aliquoted into several parts in EP tubes, incubated with apigenin under room temperature for 1 h. Samples were heated for 3 min under different temperatures (30, 35, 40, 45, 50 °C). The precipitated proteins were separated from the soluble fraction by centrifugation, and boiled at 95 °C for 10 min. Samples were subsequently used for immunoblot analysis.

### Drug affinity responsive target stabilization (DARTS) assays

DARTS assays were performed to measure the interaction stability of the small molecule apigenin with its targeting protein(s), and performed according to a previously reported protocol ^80^. PSC27 cells were lysed on ice with protease and phosphatase inhibitors, before centrifuged at 4 °C and quantified by the BCA method. Lysates were allowed to rapidly warm up to room temperature, DMSO or aigenin (100 μM) was added to the lysates for 1 h incubation in rotator. Samples were digested by pronase (1:300 (w/w), 10 μg/mL) for 30 min at room temperature. Digestion was stopped by 0.5 M EDTA (pH 8.0), with samples boiled in loading buffer for SDS-PAGE and immunoblot evaluation, or directly subjected to MS analysis for proteomic profiling. The resultant proteomic data of DARTS were deposited to the ProteomeXchange Consortium (http://proteomecentral.proteomexchange.org) through the iProX partner repository with a unique dataset identifier.

### Surface plasmon resonance (SPR) assays

The binding affinity of apigenin to recombinant human PRDX6 (rhPRDX6) was determined at 22 °C with Biacore T200 instrument equipped with CM5 sensor chips (GE healthcare) ^81^. Briefly, rhPRDX6 protein in acetate acid buffer (pH 5.5) was immobilized covalently on the sensor chips after activation by 40 mM EDC and 10 mM NHS in water solution. Different concentrations of apigenin in a gradient range of 0.78 to 12.5 μM were prepared with running buffer (phosphate-buffered saline, 0.1% sodium dodecyl sulfate and 0.05% Tween-20). The apigenin compound at different concentrations was injected simultaneously, injection of apigenin solution was performed for 60 s, with a dissociation time of 60 s in each binding cycle. Data processing and analysis were performed using Biacore 4000 and Biacore T200 evaluation software (GE healthcare).

### Immunoblot and immunofluorescence assays

Whole cell lysates were prepared using RIPA lysis buffer supplemented with protease/phosphatase inhibitor cocktail (Biomake). Nitrocellulose membranes were incubated overnight at 4 °C with primary antibodies, and HRP-conjugated goat anti-mouse or -rabbit served as secondary antibodies (Vazyme). For immunofluorescence analysis, cells were fixed with 4% formaldehyde and permeabilized before incubation with primary and secondary antibodies, each for 1 h. Upon counterstaining with DAPI (0.5 μg/mL), samples were examined with an Imager A2. Axio (Zeiss) upright microscope to analyze specific gene expression.

### Intracellular reactive oxygen species (ROS) measurements

Levels of intracellular ROS were determined using a ROS Assay Kit (Beyotime) which employs dichloro-dihydro-fluorescein diacetate (DCFH-DA) as a probe. Briefly, cells were cultured in 6-well plates for 24 h at 37 °C and were then washed twice with serum-free medium. Medium containing 10 µM DCFH-DA was added. Cells were then incubated for 20 min at 37 °C, with light avoided during incubation. After incubation, the cells were washed thrice with serum-free medium, then observed and photographed using a fluorescence microscope (Nikon). The fluorescence intensity was measured using ImageJ software (v1.51, NIH).

### *In vitro* cell phenotypic characterization

For proliferation assays of cancer cells, 2 × 10^4^ cells were dispensed into 6 well-plates and co-cultured with stromal cell-derived conditioned medium (CM). Three days later, cells were digested and counted with hemacytometer. For migration assays, cells were added to the top chambers of transwells (8 μm pore), while stromal CM were given to the bottom. Migrating cells in the bottom chambers were stained by DAPI 12-24 h later, with samples examined with an Observer A1. Axio (Zeiss) inverted microscope. Invasion assays were performed similarly with migration experiments, except that transwells were coated with basement membrane matrix (phenol red free, Corning). Alternatively, cancer cells were subject to wound healing assays performed with 6-well plates, with healing patterns graphed with a brightfield microscope. For chemoresistance assays, cancer cells were incubated with stromal CM, with the chemotherapeutic agent MIT (mitoxantrone) provided in wells for 3 days at each cell line’s IC50, a value experimentally predetermined. Cell viability was assayed by a CCK-8 kit, with the absorbance at 450 nm measured using a microplate reader.

### Histology and immunohistochemistry

Preclinical specimens from mouse tissues were fixed overnight in 10% neutral-buffered formalin and processed for paraffin embedding. Standard staining with hematoxylin/eosin was performed on sections of 5-8 μm thickness cut from each specimen block. For immunohistochemistry, tissue sections were de-paraffinized and incubated in citrate buffer at 95 °C for 40 min for antigen retrieval before incubated with primary antibodies (*e.g*., anti-cleaved Caspase 3, 1:1000) overnight at 4°C. After 3 washes with PBS, tissue sections were incubated with biotinylated secondary antibody (1:200 dilution, Vector Laboratories) for 1 h at room temperature then washed thrice, after which streptavidin-horseradish peroxidase conjugates (Vector Laboratories) were added and the slides incubated for 45 min. DAB solution (Vector Laboratories) was then added, with slides counterstained with haematoxylin.

### Experimental animals and chemotherapeutic studies

Animals were maintained in a specific pathogen-free (SPF) facility, with NOD/SCID (Model Animal Research Center of Nanjing University) mice at an age of approximately 6 weeks (∼20 g body weight) used. Ten mice were incorporated in each group, with xenografts subcutaneously generated at the hind flank upon anesthesia mediated by isoflurane inhalation. Stromal cells (PSC27) were mixed with cancer cells (PC3) at a ratio of 1:4 (i.e., 250,000 stromal cells admixed with 1,000,000 cancer cells to make tissue recombinants before implantation *in vivo*).

Animals were sacrificed at 2-8 weeks after tumor implantation, according to tumor burden or experimental requirements. Tumor growth was monitored weekly, with tumor volume (v) measured and calculated according to the tumor length (l), width (w) and height (h) by the formula: v = (π/6) × ((l+w+h)/3)^3 82^. Freshly dissected tumors were either snap-frozen or fixed to prepare FFPE samples. Resulting sections were used for IHC staining against specific antigens or subject to hematoxylin/eosin staining.

For chemoresistance studies, animals received subcutaneous implantation of tissue recombinants as described above and were given standard laboratory diets for 2 weeks to allow tumor uptake and growth initiation. Starting from the 3^rd^ week (tumors reaching 4-8 mm in diameter), MIT (0.2 mg/kg doses), the senomorphic agent apigenin (10.0 mg/kg doses, 200 μl/dose) or vehicle controls was administered through intraperitoneal injection (therapeutic agents *via* i.p. route), on the 1^st^ day of 3^rd^, 5^th^ and 7^th^ weeks, respectively. Upon completion of the 8-week therapeutic regimen, animals were sacrificed, with tumor volumes recorded and tissues processed for histological evaluation.

At the end of chemotherapy and/or targeting treatment, animals were anaesthetized and peripheral blood was gathered *via* cardiac puncture. Blood was transferred into a 1.5 ml Eppendorf tube and kept on ice for 45 min, followed by centrifugation at 9000 x g for 10 min at 4 °C. Clear supernatants containing serum were collected and transferred into a sterile 1.5 ml Eppendorf tube. All serum markers were measured using dry-slide technology on IDEXX VetTest 8008 chemistry analyzer (IDEXX). About 50 μl of the serum sample was loaded on the VetTest pipette tip before securely fit on the pipettor and manufacturer’s instructions were followed for further examination.

All animal experiments were performed in compliance with NIH Guide for the Care and Use of Laboratory Animals (National Academies Press, 2011) and the ARRIVE guidelines, and were approved by the Institutional Animal Care and Use Committee (IACUC) of Shanghai Institute of Nutrition and Health, Chinese Academy of Sciences.

### Tissue SA-β-Gal staining and histological examination

For SA-β-Gal staining, frozen sections were dried at 37 °C for 20-30 min before fixed for 15 min at room temperature. The frozen sections were washed thrice with PBS and incubated with SA-β-Gal staining reagent (Beyotime) overnight at 37 °C. After completion of SA-β-Gal staining, sections were stained with eosin for 1-2 min, rinsed under running water for 1 min, differentiated in 1% acid alcohol for 10-20 s, and washed again under running water for 1 min. Sections were dehydrated in increasing concentrations of alcohol and cleared in xylene. After drying, samples were examined under a bright-field microscope.

### Tissue immunostaining analyses

For immunohistochemistry, tumor specimens were fixed in 4% paraformaldehyde and embedded in paraffin. Sections of 5 μm thick were stained with antibody against IL6 or cleaved caspase 3 (CCL3) for SASP expression appraisal and cell apoptosis evaluation, respectively. Briefly, heat-induced antigen retrieval was performed using 10 mM sodium citrate buffer, pH 6.5. Peroxidase activity was quenched with 3% H_2_O_2_ and tissues were blocked in 5% bovine serum albumin for 30 min. Primary antibodies (1:100-1:200 dilution) were incubated on sections for overnight at 4 °C. Horseradish peroxidase-conjugated secondary antibody (1:1000) and DAB were used for detection. Slides were counterstained with hematoxylin. Alternatively, frozen tissue sections were used for immunofluorescence staining of p21 induction in animals receiving various preclinical treatments, with counterstaining performed with DAPI. Images were captured with epifluorescence microscope.

### Appraisal of *in vivo* cytotoxicity by blood tests

For routine blood examination, 100 μl fresh blood was acquired from each animal and mixed with EDTA immediately. The blood samples were analyzed with Celltac Alpha MEK-6400 series hematology analyzers (Nihon Kohden). For serum biochemical analyses, blood samples were collected and clotted for 2 h at room temperature or overnight at 4 °C. Samples were then centrifuged (1000 × g, 10 min) to obtain serum. An aliquot of approximately 50 μl serum was subject to analysis for creatinine, urea, alkaline phosphatase (ALP) and alanine transaminase (ALT) by an automatic biochemical analyzer (BS-5800M, Mindray Bio-Medical Electronics Co. Ltd). Evaluation of circulating levels of hemoglobin, white blood cells, lymphocytes and platelets were performed using dry-slide technology on a VetTest 8008 chemistry analyzer (IDEXX) as reported previously ^82^.

All animal experiments were conducted in compliance with the NIH Guide for the Care and Use of Laboratory Animals (National Academies Press, 2011) and the ARRIVE guidelines, and were approved by the IACUC of Shanghai Institute of Nutrition and Health, Chinese Academy of Sciences. For each preclinical regimen, animals were monitored for conditions including hypersensitivity (changes in body temperature, altered breathing and ruffled fur), body weight, mortality and changes in behavior (*i.e.*, loss of appetite and distress), and were disposed of appropriately according to the individual pathological severity as defined by relevant guidelines.

### Open field test

The open-field test was employed to assess the locomotor activity and anxiety of experimental animals when placed in an unfamiliar environment ^83^. The open-field chambers (l × w × h, 60 cm × 60 cm × 60 cm) were made of Plexiglas with a non-reflective square base. Total distance in the whole open-field chamber, distance traveled in the center zone (30 cm × 30 cm, 50% of the total area) and in the edge zone (20% of the total area, along edges) were recorded. Less activity in the center zone indicated anxiety-like behaviors.

### Spontaneous alternation Y Maze test

Spontaneous alternation behavior indicates the tendency for mice to alternate their (conventionally) nonreinforced choices on successive opportunities ^84^. The Y maze spontaneous alternation was employed to assess short-time spatial recognition and working memory in mice by measurement of spontaneous alternations ^85^. The maze consisted of three arms, with each arm 31 cm long, 5 cm wide and 10 cm high. Each arm had markers of different colors as distinct visual cues. After placed individually at the center of the apparatus, mice were allowed to explore freely through the maze during an 8-min session. Alternation was defined as successive entries into all three arms on overlapping triplet sets. The number of arm entries and alternations were recorded visually to calculate the percentage of the alternation behavior with the following formula: % Alternation = [Number of Alternations/(Total number of arm entries -2)] x 100. Spontaneous alternation (%), defined as successive entries into the three arms on overlapping triplet sets, is associated with spatial short-term memory.

### Statistical analysis

All *in vitro* experiments were performed in triplicates, while animal studies were conducted with at least 8 mice *per* group for most preclinical assays. Data are presented as mean ± SD except where otherwise indicated. GraphPad Prism 9.5.1 was used to collect and analyze data, with statistical significance determined according to individual settings. Cox proportional hazards regression model and multivariate Cox proportional hazards model analyses were performed with statistical software SPSS. Statistical significance was determined by unpaired two-tailed Student’s *t* tests, one- or two-way ANOVA, Pearson’s correlation coefficients tests, Kruskal-Wallis, log-rank tests, Wilcoxon-Mann-Whitney tests or Fisher’s exact tests. For all statistical tests, a *P* value < 0.05 was considered significant.

### Data availability

Experimental data supporting the plots within this paper and other findings of this study are available from the corresponding author upon reasonable request. The RNA-seq data generated in the present study have been deposited in the Gene Expression Omnibus database under accession code GSE273159.

