## Supplemental File(s) for "Repurposing the plant-derived compound apigenin for senomorphic effect in antiaging pipelines"

#### Figure S1

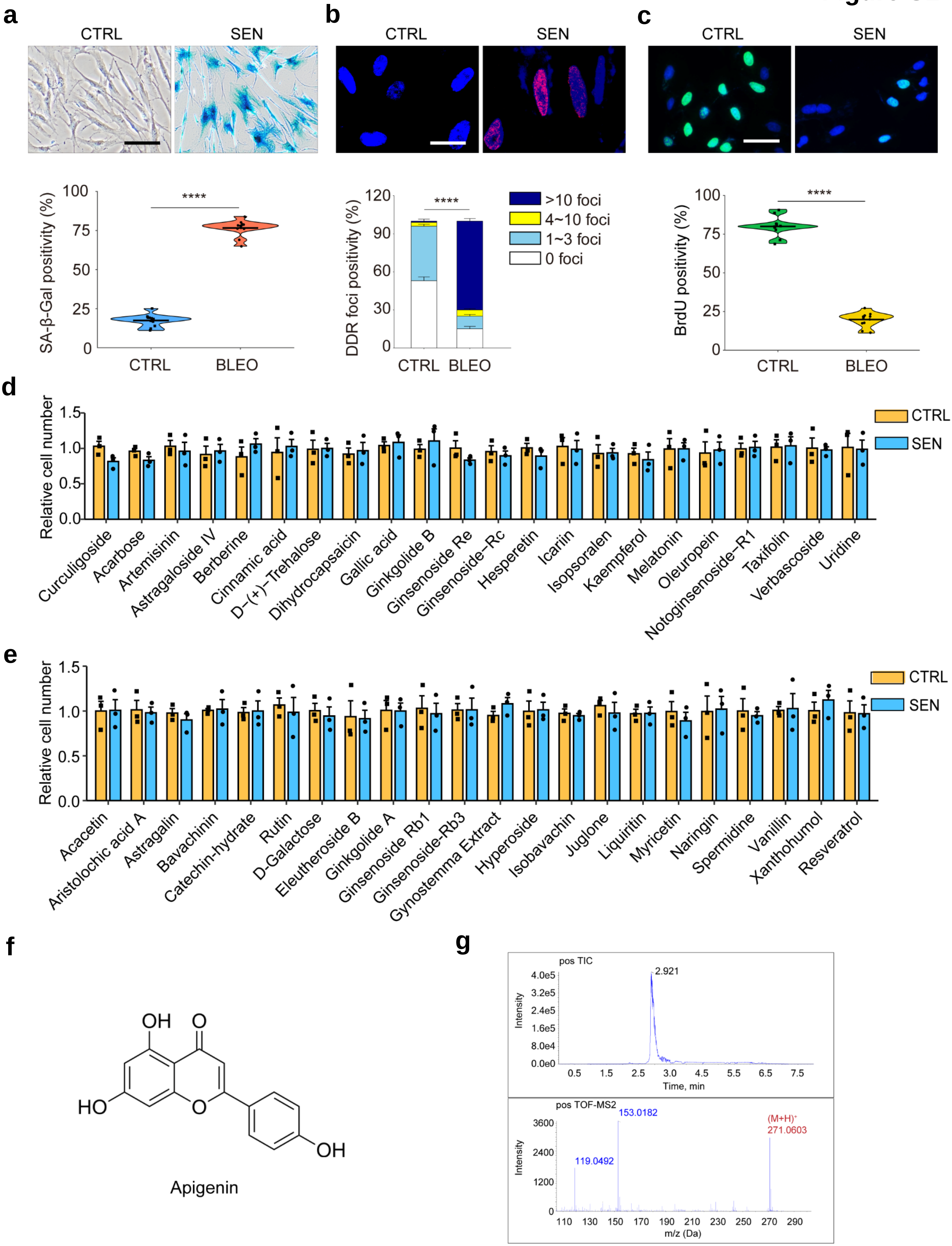

a
CTRL
SEN
SA-β-Gal positivity (%)
CTRL
BLEO
****
b
CTRL
SEN
DDR foci positivity (%)
>10 foci
4~10 foci
1~3 foci
0 foci
c
CTRL
SEN
BrdU positivity (%)
d
Relative cell number
CTRL
SEN
Curculigoside
Acarbose
Artemisinin
Astragaloside IV
Berberine
Cinnamic acid
D-(+)-Trehalose
Dihydrocapsaicin
Gallic acid
Ginkgolide B
Ginsenoside Re
Ginsenoside-Rc
Hesperetin
Icariin
Isopsoralen
Kaempferol
Melatonin
Oleuropein
Notoginsenoside-R1
Taxifolin
Verbascoside
Uridine
e
Acacetin
Aristolochic acid A
Astragalin
Bavachinin
Catechin-hydrate
Rutin
D-Galactose
Eleutheroside B
Ginkgolide A
Ginsenoside Rb1
Ginsenoside-Rb3
Gynostemma Extract
Hyperoside
Isobavachin
Juglone
Liquiritin
Myricetin
Naringin
Spermidine
Vanillin
Xanthohumol
Resveratrol
f
Apigenin
g
pos TIC
2.921
Intensity
Time, min
pos TOF-MS2
153.0182
119.0492
(M+H)+
271.0603
m/z (Da)

### Figure S2

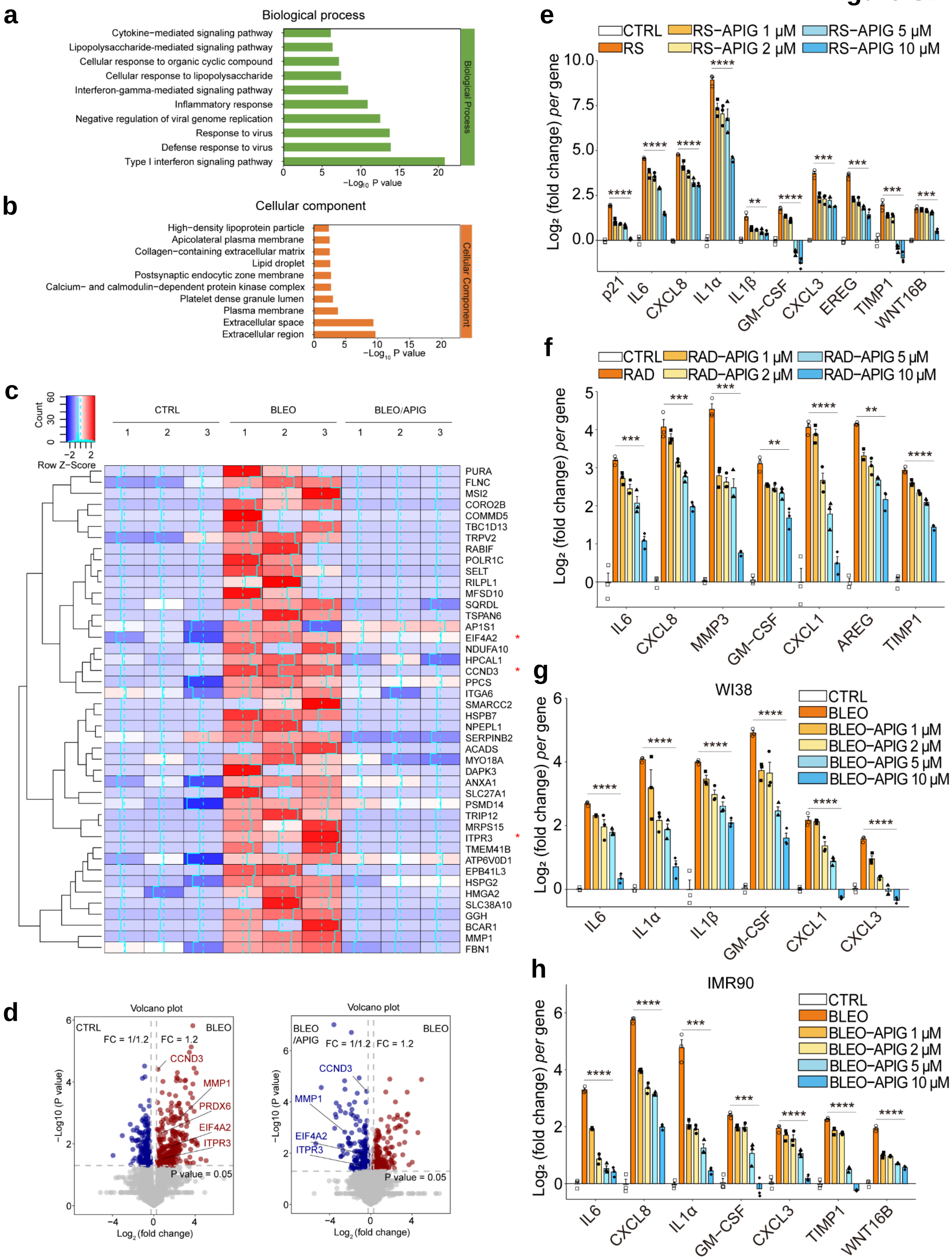

a
Biological process
Cytokine−mediated signaling pathway
Lipopolysaccharide−mediated signaling pathway
Cellular response to organic cyclic compound
Cellular response to lipopolysaccharide
Interferon−gamma−mediated signaling pathway
Inflammatory response
Negative regulation of viral genome replication
Response to virus
Defense response to virus
Type I interferon signaling pathway
Biological Process
−Log10 P value
b
Cellular component
High−density lipoprotein particle
Apicolateral plasma membrane
Collagen−containing extracellular matrix
Lipid droplet
Postsynaptic endocytic zone membrane
Calcium− and calmodulin−dependent protein kinase complex
Platelet dense granule lumen
Plasma membrane
Extracellular space
Extracellular region
Cellular Component
−Log10 P value
c
Count
Row Z−Score
CTRL
BLEO
BLEO/APIG
PURA
FLNC
MSI2
CORO2B
COMMD5
TBC1D13
TRPV2
RABIF
POLR1C
SELT
RILPL1
MFSD10
SQRDL
TSPAN6
AP1S1
EIF4A2
NDUFA10
HPCAL1
CCND3
PPCS
ITGA6
SMARCC2
HSPB7
NPEPL1
SERPINB2
ACADS
MYO18A
DAPK3
ANXA1
SLC27A1
PSMD14
TRIP12
MRPS15
ITPR3
TMEM41B
ATP6V0D1
EPB41L3
HSPG2
HMGA2
SLC38A10
GGH
BCAR1
MMP1
FBN1
d
Volcano plot
CTRL
BLEO
FC = 1/1.2
FC = 1.2
CCND3
MMP1
PRDX6
EIF4A2
ITPR3
P value = 0.05
−Log10 (P value)
Log2 (fold change)
Volcano plot
BLEO /APIG
BLEO
FC = 1/1.2
FC = 1.2
CCND3
MMP1
EIF4A2
ITPR3
P value = 0.05
−Log10 (P value)
Log2 (fold change)
e
CTRL
RS
RS−APIG 1 μM
RS−APIG 2 μM
RS−APIG 5 μM
RS−APIG 10 μM
Log2 (fold change) per gene
p21
IL6
CXCL8
IL1α
IL1β
GM−CSF
CXCL3
EREG
TIMP1
WNT16B
f
CTRL
RAD
RAD−APIG 1 μM
RAD−APIG 2 μM
RAD−APIG 5 μM
RAD−APIG 10 μM
Log2 (fold change) per gene
IL6
CXCL8
MMP3
GM−CSF
CXCL1
AREG
TIMP1
g
WI38
CTRL
BLEO
BLEO−APIG 1 μM
BLEO−APIG 2 μM
BLEO−APIG 5 μM
BLEO−APIG 10 μM
Log2 (fold change) per gene
IL6
IL1α
IL1β
GM−CSF
CXCL1
CXCL3
h
IMR90
CTRL
BLEO
BLEO−APIG 1 μM
BLEO−APIG 2 μM
BLEO−APIG 5 μM
BLEO−APIG 10 μM
Log2 (fold change) per gene
IL6
CXCL8
IL1α
GM−CSF
CXCL3
TIMP1
WNT16B

**Figure S3**

**a**

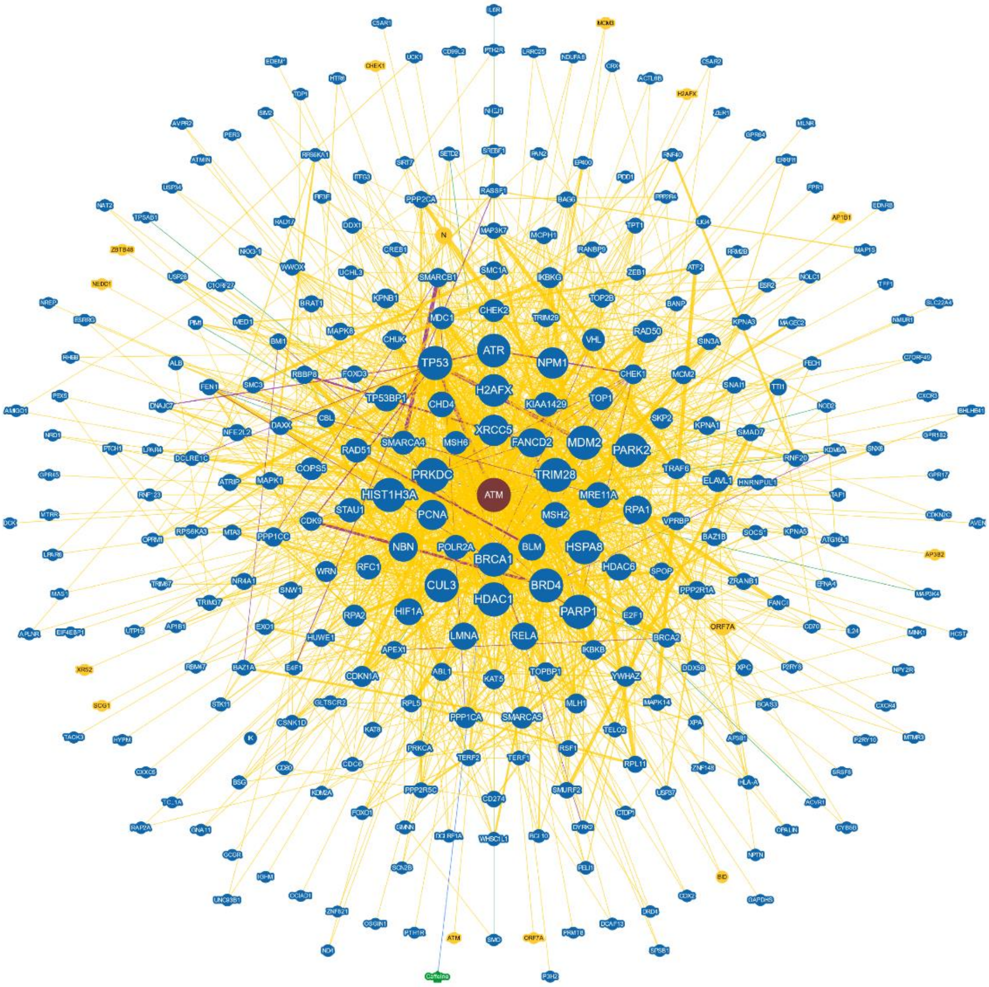
ATM
TP53
ATR
NPM1
H2AFX
TP53BP1
CHD4
XRCC5
KIAA1429
MDM2
PARK2
FANCD2
PRKDC
TRIM28
HIST1H3A
PCNA
NBN
BRCA1
CUL3
HDAC1
BRD4
PARP1
HSPA8
BLM
MSH2
RPA1
MRE11A
LMNA
RELA
HIF1A

**b**

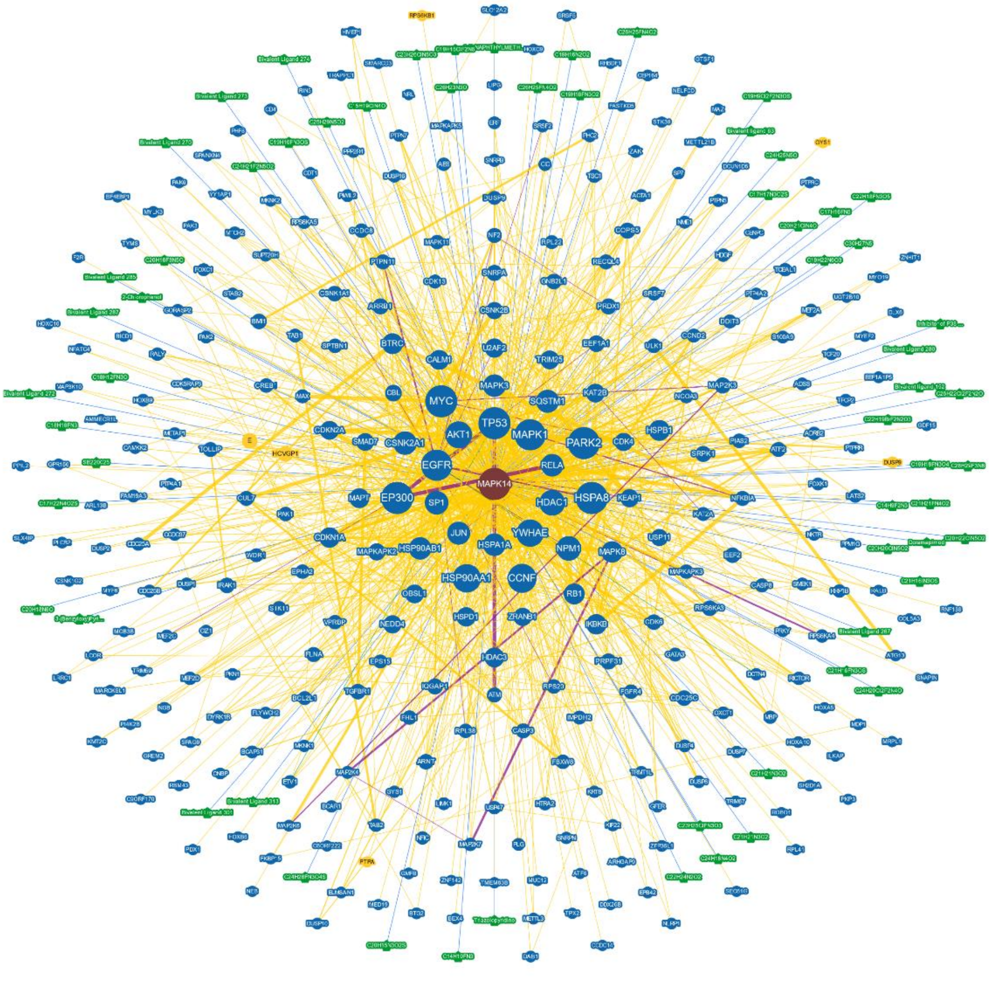
MAPK14
MYC
TP53
AKT1
MAPK1
PARK2
EGFR
EP300
SP1
HSPA8
HDAC1
YWHAE
JUN
HSP90AA1
HSP90AB1
CCNF
NPM1
HSPA1A

**Figure S4**

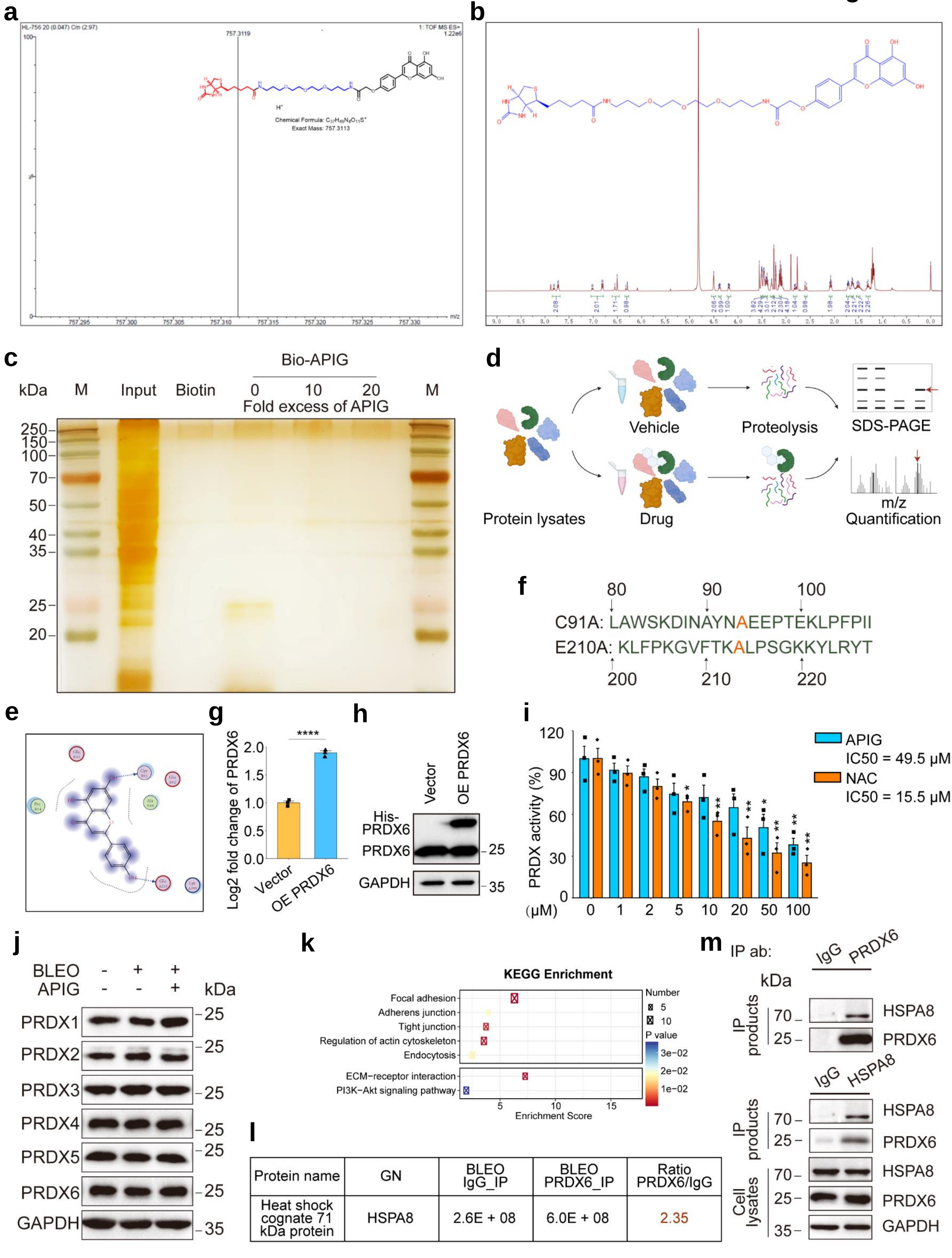


| Protein name | GN | BLEO IgG_IP | BLEO PRDX6_IP | Ratio PRDX6/IgG |
|---|---|---|---|---|
| Heat shock cognate 71 kDa protein | HSPA8 | 2.6E + 08 | 6.0E + 08 | 2.35 |

### Figure S5

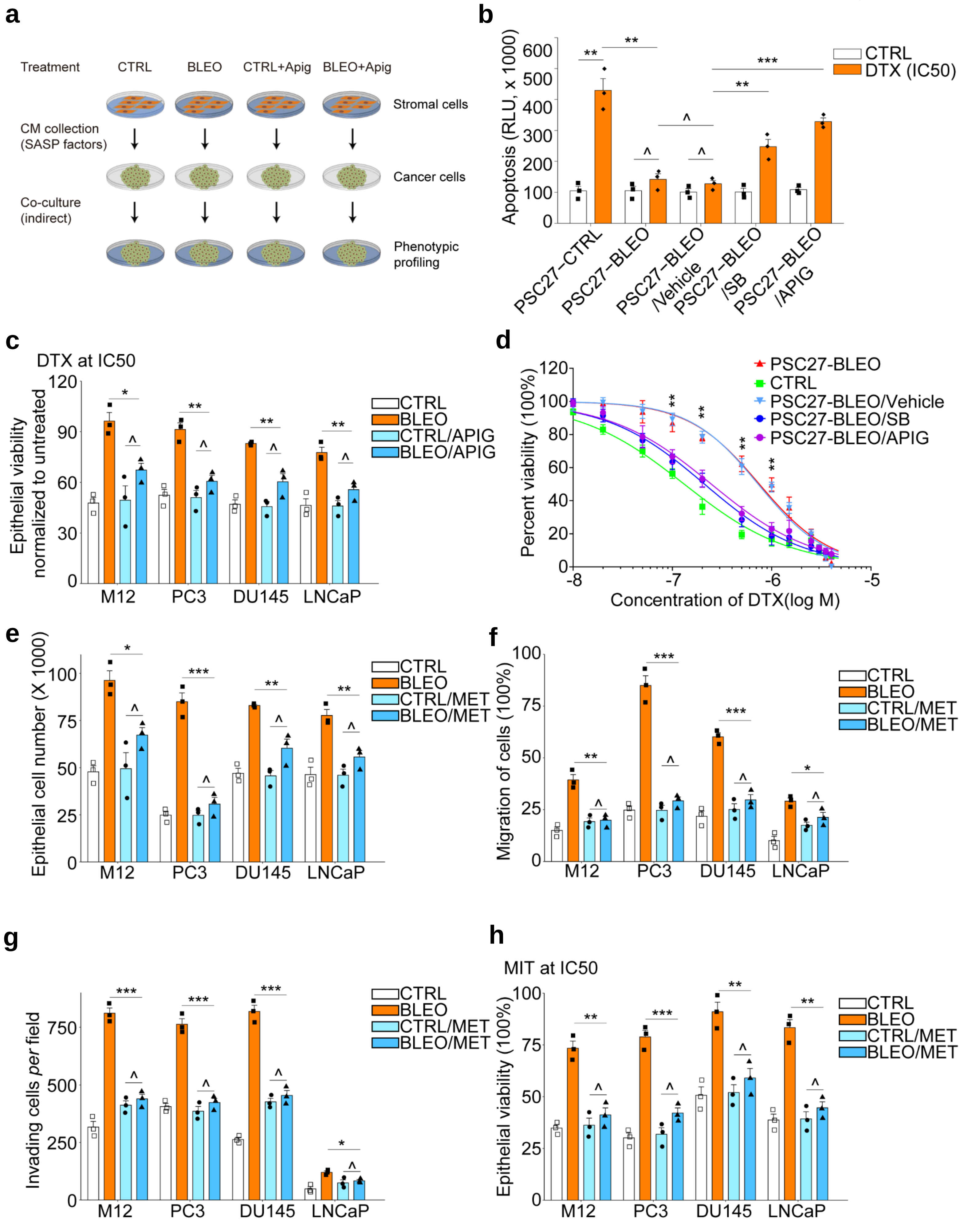
a
Treatment
CTRL
BLEO
CTRL+Apig
BLEO+Apig
Stromal cells
CM collection
(SASP factors)
Cancer cells
Co-culture
(indirect)
Phenotypic
profiling
b
Apoptosis (RLU, x 1000)
CTRL
DTX (IC50)
PSC27–CTRL
PSC27–BLEO
PSC27–BLEO
/Vehicle
PSC27–BLEO
/SB
PSC27–BLEO
/APIG
c
DTX at IC50
Epithelial viability
normalized to untreated
CTRL
BLEO
CTRL/APIG
BLEO/APIG
M12
PC3
DU145
LNCaP
d
Percent viability (100%)
PSC27-BLEO
CTRL
PSC27-BLEO/Vehicle
PSC27-BLEO/SB
PSC27-BLEO/APIG
Concentration of DTX(log M)
e
Epithelial cell number (X 1000)
CTRL
BLEO
CTRL/MET
BLEO/MET
M12
PC3
DU145
LNCaP
f
Migration of cells (100%)
CTRL
BLEO
CTRL/MET
BLEO/MET
M12
PC3
DU145
LNCaP
g
Invading cells per field
CTRL
BLEO
CTRL/MET
BLEO/MET
M12
PC3
DU145
LNCaP
h
MIT at IC50
Epithelial viability (100%)
CTRL
BLEO
CTRL/MET
BLEO/MET
M12
PC3
DU145
LNCaP

Figure S6

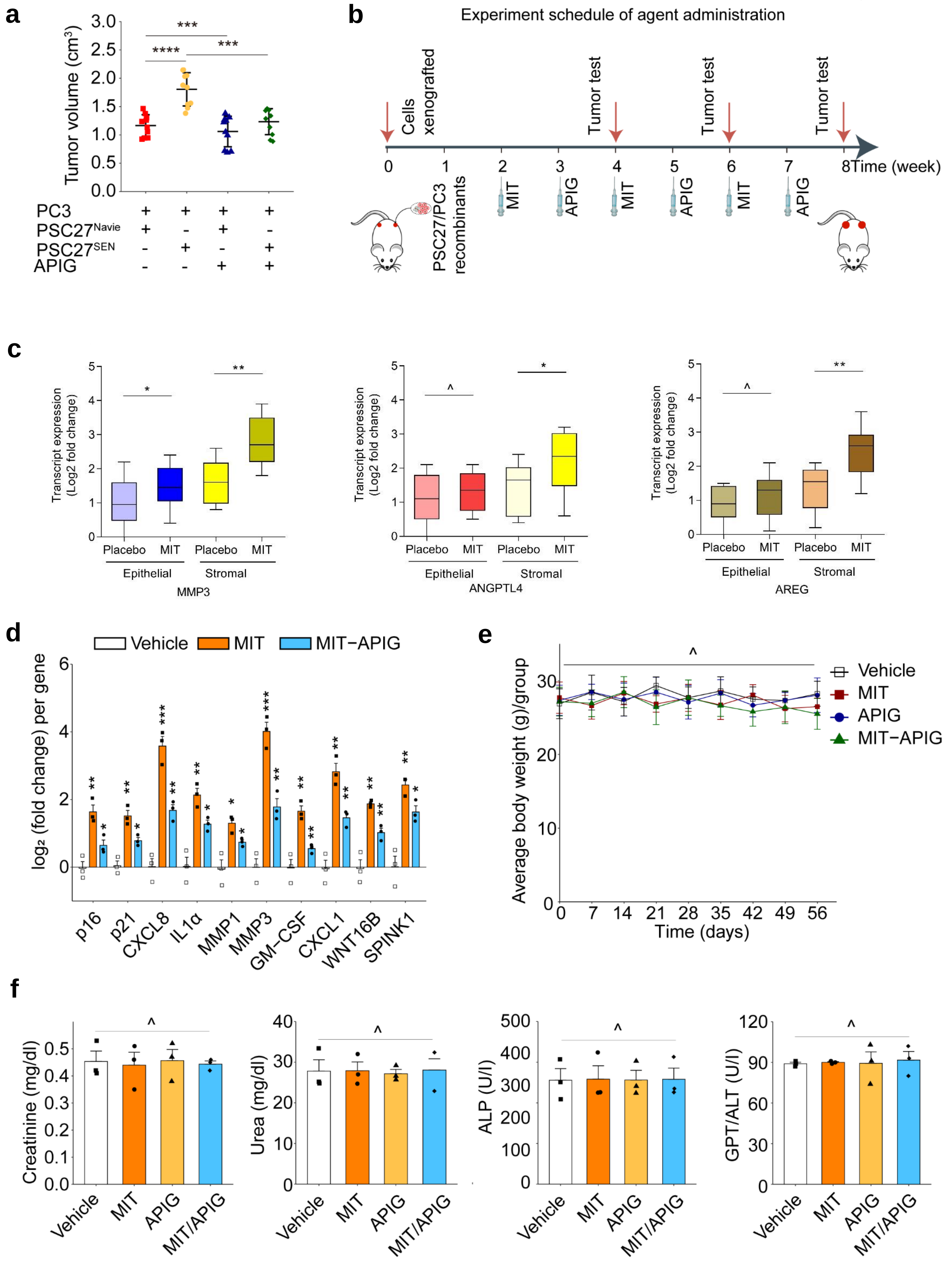

a
Tumor volume (cm3)
PC3
PSC27Navie
PSC27SEN
APIG
b
Experiment schedule of agent administration
Cells xenografted
Tumor test
Time (week)
PSC27/PC3 recombinants
MIT
APIG
c
Transcript expression (Log2 fold change)
Placebo
MIT
Epithelial
Stromal
MMP3
ANGPTL4
AREG
d
Vehicle
MIT
MIT−APIG
log2 (fold change) per gene
p16
p21
CXCL8
IL1α
MMP1
MMP3
GM-CSF
CXCL1
WNT16B
SPINK1
e
Average body weight (g)/group
Time (days)
Vehicle
MIT
APIG
MIT−APIG
f
Creatinine (mg/dl)
Urea (mg/dl)
ALP (U/l)
GPT/ALT (U/l)
Vehicle
MIT
APIG
MIT/APIG

Figure S7

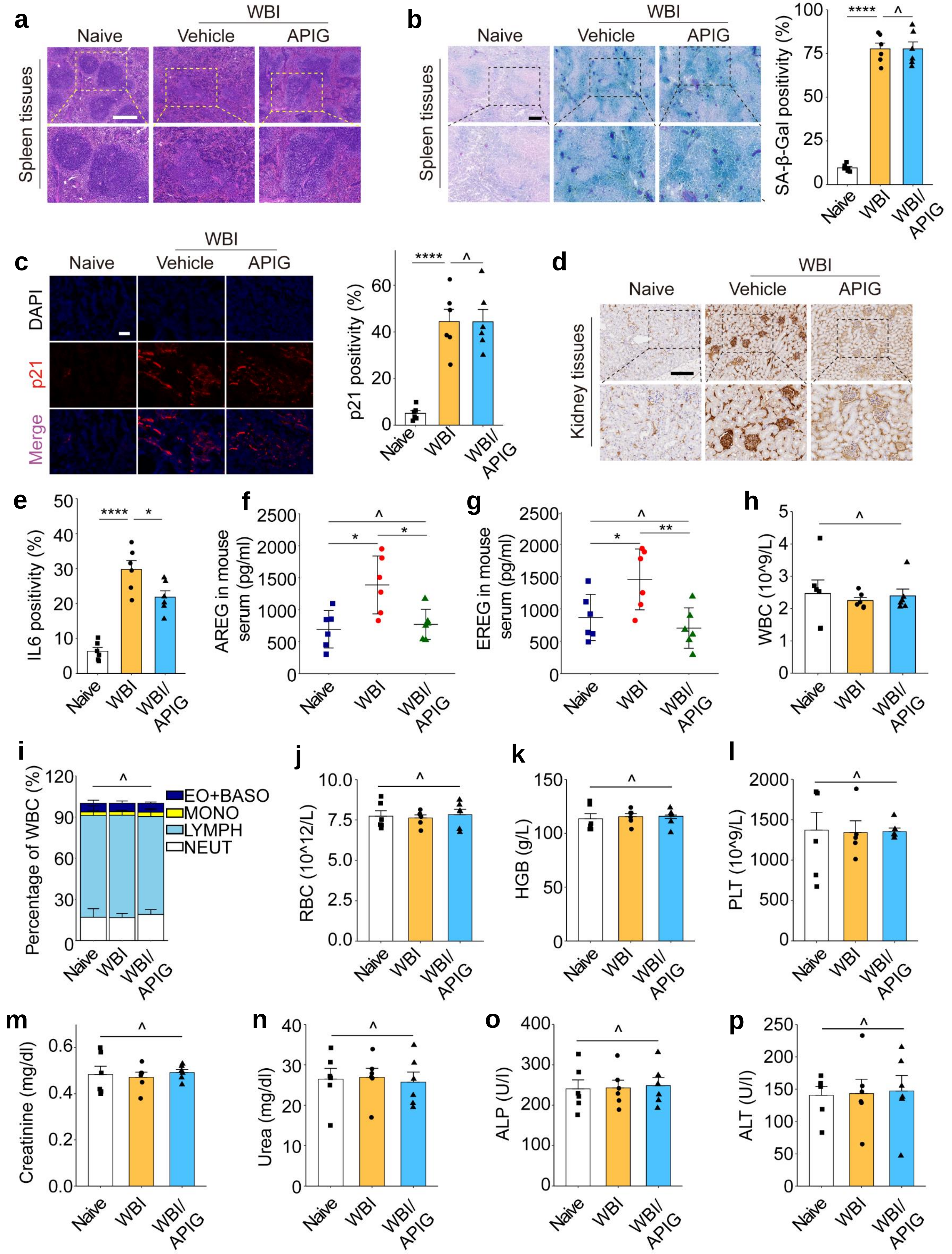
a
WBI
Naive
Vehicle
APIG
Spleen tissues
b
WBI
Naive
Vehicle
APIG
Spleen tissues
SA-β-Gal positivity (%)
****
^
Naive
WBI
WBI/APIG
c
WBI
Naive
Vehicle
APIG
DAPI
p21
Merge
p21 positivity (%)
****
^
d
WBI
Naive
Vehicle
APIG
Kidney tissues
e
IL6 positivity (%)
****
*
f
AREG in mouse serum (pg/ml)
^
*
*
g
EREG in mouse serum (pg/ml)
^
*
**
h
WBC (10^9/L)
^
i
Percentage of WBC (%)
^
EO+BASO
MONO
LYMPH
NEUT
j
RBC (10^12/L)
^
k
HGB (g/L)
^
l
PLT (10^9/L)
^
m
Creatinine (mg/dl)
^
n
Urea (mg/dl)
^
o
ALP (U/l)
^
p
ALT (U/l)
^

**Supplementary Fig. 1 Phenotypic profiling of TIS, senolytics screening and a potential senomorphic candidate apigenin.** (a) Representative images of senescence appraisal by SA-β-Gal staining of PSC27 cells upon occurrence of TIS. Scale bar, 15 μm. Lower, statistics. (b) Representative images of BrdU staining to assess DNA incorporation of PSC27 cells. Scale bar, 5 μm. Lower, statistics. (c) Representative images of DDR by immunofluorescence staining of γh2AX in PSC27 cells. The DDR profile was categorized into 4 sub-groups including 0 foci, 1-3 foci, 4-10 foci and >10 foci *per* cell. Scale bar, 10 μm. Lower, statistics. (d-e) Evaluation of the senolytic potential of remaining natural products (10 μM/agent) in SEN and CTRL cells. (f) Chemical structure of the natural flavonoid apigenin. (g) Mass spectrometry plots displaying base peak chromatogram spectra, precursor ion (MS1) and fragment ion (MS2) of apigenin by performing HPLC-QTOF-MS/MS. Data in a and c are plotted as violin graphs and representative of 10 independent biological replicates with *P* values calculated by Student's *t* tests. TIS, therapy-induced senescence. DDR, DNA damage response. Data in b, d and e are shown as mean ± SD and are representative of 3 independent biological replicates with *P* values calculated by two-way ANOVA with Turkey's multiple-comparison tests (b) or Student's *t* tests (d, e). ^, $P > 0.05$; *, $P < 0.05$; **, $P < 0.01$; ***, $P < 0.001$; ****, $P < 0.0001$.

**Supplementary Fig. 2 Transcriptome- and proteome-wide expression analysis of senescent cells upon apigenin treatment.** (a-b) Bar plots showing GO aspects including biological process (BP, a) and cellular component (CC, b) of 67 human genes most significantly restrained by apigenin in senescent PSC27 cells. (c) Heatmap profiling of proteomic expression in control cells, senescent cells and senescent cells exposed to apigenin (10 μM) as examined by mass spectrometry. Red stars, senescence-associated proteins. (d) Volcano plot displaying the differentially expressed proteins (red, upregulated; blue, downregulated) in control cells, senescent cells or senescent cells treated by apigenin. (e) Quantitative appraisal of the expression of canonical SASP factors at transcriptional level upon replicative senescence (RS) at increasing apigenin concentrations. (f) Quantitative evaluation of canonical SASP factor expression at transcriptional level upon ionizing radiation (RAD)-induced senescence with cells exposed to increasing apigenin concentrations. (g) Quantitative analysis of canonical SASP factor expression at transcriptional level in human lung fibroblast cell line WI38 upon BLEO-induced senescence with cells exposed to increasing apigenin concentrations. (h) Quantitative evaluation of canonical SASP factor expression at transcriptional level in human lung fibroblast cell line IMR90 upon BLEO-induced senescence with cells exposed to increasing apigenin concentrations. Unless specially noted, data in e-h are shown as mean ± SD and representative of 3 independent biological replicates with $P$ values calculated by Student's $t$ tests. ^, $P > 0.05$; ^, $P > 0.05$; *, $P < 0.05$; **, $P < 0.01$; ***, $P < 0.001$; ****, $P < 0.0001$.

**Supplementary Fig. 3 Proteome-wide mapping of potential protein-protein interactions of target molecules.** (a) Mapping of the interactive proteins of ATM with BioGRID, a biomedical interaction repository with data compiled through comprehensive curation and serving as an open access database archiving and sharing protein interaction data from model organism species and humans. (b) A BioGRID-based mapping of p38 MAPK-interactive molecules, with the biomedical interaction repository as described in (a).

**Supplementary Fig. 4 The interaction of apigenin with PRDX6 results in suppressed iPLA2 activity.** (a) High resolution MS of Bio-APIG probe, which connects biotin and hydroxyl of apigenin with bio-inert PEG). (b) $^{1}$H NMR spectrum of biotin-apigenin probe (400 MHz, methanol-d4). (c) Lysates of senescent PSC27 cells were treated by Bio-APIG or biotin with or without a 10- or 20-fold excess of unlabeled apigenin, before subjected to pull-down by streptavidin-agarose. Proteins captured in beads were determined by SDS-PAGE and subject to silver staining. (d) A schematic illustration introducing the process of DARTS, a label-free drug target discovery approach. PSC27 cell lysates were incubated with apigenin or vehicle for

1 h, followed by proteolysis with pronase for 0.5 h and MS for differential protein quantification. (e) Two dimensional *in silico* molecular modelling of apigenin potentially bound to Cys91 and Glu210 of PRDX6 as depicted in Fig 4k. (f) Diagram showing peptide sequences of PRDX6 with two amino acid mutations in the active site. (g-h) HEK293T cells were transfected with a PRDX6 expression construct encoding His-tagged PRDX6 sequence for stable overexpression. (g) Quantitative analysis of PRDX2 expression in cells transfected with the His-tagged PRDX6 construct or vector control. (h) Immunoblot examination of protein expression in cells as described in (g). GAPDH, loading control. (i) Examination of peroxidase activity by measuring the remaining $H_2O_2$ levels in HEK293T cells stably overexpressing PRDX6 after incubation with gradient concentrations of apigenin or NAC, a peroxide scavenger, for 30 min. (j) Immunoblot profiling of the PRDX family (PRDX1-6) at protein level in stromal cells exposed to BLEO and/or apigenin. GAPDH, loading control. (k) KEGG pathway enrichment analysis of significantly downregulated proteins at proteomic level in senescent cells treated with MJ33, a selective PRDX6-iPLA2 inhibitor, as compared with cells treated with vehicle. (l-m) Immunoprecipitation (IP) coupled with MS analysis or immunoblot assay to detect protein-protein interactions between PRDX6 and HSPA8. (l) Senescent cells were lysed for IP with IgG or anti-PRDX6, with the enriched beads analyzed by MS thereafter. (m) Senescent cells were lysed for IP with IgG, anti-PRDX6 or anti-HSPA8, with both immunoprecipitates (IPs) and inputs analyzed. MS, mass spectrometry. Bio-APIG, biotin-apigenin. DARTS, drug affinity responsive target stability. Data in g and k are shown as mean ± SD and representative of 3 independent biological replicates, with *P* values calculated by Student's *t*-tests. ^, $P > 0.05$; ^, $P > 0.05$; *, $P < 0.05$; **, $P < 0.01$; ***, $P < 0.001$; ****, $P < 0.0001$.

**Supplementary Fig. 5 Apigenin reduces PCa cell malignancy conferred by senescent stromal cell-derived conditioned media.** (a) A schematic workflow showing the procedure of the indirect co-culture involving stromal cells and PCa cells. CM containing a myriad of soluble factors were collected from native or senescent stromal cells 8-10 d post BLEO treatment, with cellscultured in the absence or presence of apigenin. PCa cells including PC3, DU145, M12 and LNCaP were examined for phenotypic assessments. The CM of an equal number of cells was collected under each condition. (b) Apoptotic activity measurement of PCa cells in culture supplemented with half-maximal inhibitory concentration (IC50) of DTX. Signal readings proportional to the caspase-3/7 activity of PC3 cells were depicted by relative luminescence units (RLUs). (c) Chemoresistance assay of PCa cells exposed to DTX while being cultured with different types of CM. DTX was administered at a pre-determined IC50 for each cell line. (d) Dose-response curves (nonlinear regression/curve fit) of PC3 cells cultured with the different types of CM, and concurrently exposed to a wide range of concentrations of DTX. Data were plotted on an exponential scale, with cell viability examined relative to the untreated group and calculated as a percentage. (e) Proliferation assay of PCa cells, which were incubated for 3 days with different types of CM as indicated in (a). (f) Migration assay of PCa cells incubated with several types of CM for 18 h. (g) Invasiveness assay of PCa cells across collagen-based transwell membrane incubated with different types of CM. (h) Chemoresistance assay of PCa cells treated with MIT while being cultured with different types of CM. MIT was added at a pre-determined IC50 for each cell line. In e-h, metformin (MET) was used instead of apigenin. PCa, prostate cancer. CM, conditioned media. DTX, docetaxel (a chemotherapeutic agent). MIT, mitoxantrone. SB, SB203580 (a p38MAPK inhibitor that can block the SASP expression). Data in b, c, d, e, f, g and h are shown as mean ± SD and representative of 3 independent biological replicates, with *P* values calculated by Student's *t*-tests. ^, $P > 0.05$; ^, $P > 0.05$; *, $P < 0.05$; **, $P < 0.01$; ***, $P < 0.001$; ****, $P < 0.0001$.

**Supplementary Fig. 6 Experimental design of the preclinical regimen, stroma-conferred therapeutic resistance and appraisal of tumor regression in mice receiving administration of MIT and/or apigenin.** (a) Comparative statistics of tumor volumes at the end of an 8-week period in NOD/SCID mice carrying PC3 cells alone or admixed with PSC27 cells in the hind flank. PSC27 cells were either naive or senescent induced by BLEO prior to inoculation ($PSC27^{Naive}$ and $PSC27^{SEN}$, respectively). (b) Schematic diagram of drug administration and tumor evaluation of the preclinical trial. Two weeks prior to chemotherapy, PC3 cells alone or together with PSC27 cells were xenografted subcutaneously to animals. MIT was administrated *via* i.v. on the 1st day of each week starting from the 3rd week, then delivered every other week with a total number of 3 doses. Apigenin was delivered *via* i.p. on the 1st day of every other week starting from $5^{th}$ week. Mice were sacrificed at the end of the 8-week regimen for tumor evaluation and histological assessment. (c) Transcriptional evaluation of several typical SASP factors (MMP3, ANGPTL4 and AREG) in stromal cells isolated from PC3/PSC27 tumor foci *via* laser capture microdissection (LCM). (d) Transcriptional analysis of several typical SASP factors in stromal cells as depicted in (c). Signals were normalized to vehicle-treated group *per* gene. (e) Measurement of mouse body weights once a week throughout the whole period of the therapeutic regimen. (f) Terminal bleeds were taken from retro-orbital at the end of therapeutic regimens, and subject to analysis of circulating concentrations of creatinine, urea, alkaline phosphatase (ALP) and alanine aminotransferase (ALT) for toxicity evaluation. Data in c, d, e and f are shown as mean ± SD and representative of 3 independent biological replicates, with *P* values calculated by Student's *t*-tests. ^, P > 0.05; ^, $P > 0.05$; *, $P < 0.05$; **, $P < 0.01$; ***, $P < 0.001$; ****, $P < 0.0001$.

**Supplementary Fig. 7 Apigenin treatment alleviates physical frailty and reduces pathological indices of animals prematurely aged after WBI.** (a) H&E staining to histologically profile the morphological changes of spleen tissues of C57BL/6J mice. Naïve animals, WBI-treated animals experiencing vehicle treatment and WBI-treated animals receiving apigenin administration, respectively, were recruited to the preclinical study. Scale bar, 400 μm. (b) Representative images (left) and comparative quantification (right) of SA-β-Gal staining of spleen tissues dissected from experimental mice. Animals experiencing different treatments as described in (a) were assessed. Scale bar, 100 μm. (c) IF staining of p21 expressed in kidney tissues. Scale bar, 20 μm. Left, representative images. Right, comparative statistics. (d) Representative images of IHC staining to detect IL6 expression in kidney tissues. Scale bar, 200 μm. (e) Statistic comparison of IL6 staining positivity in kidney tissues as examined in (d). (f-g) ELISA measurement of circulating concentrations of typical SASP factors represented by AREG (f) and EREG (g) in animal serum. (h) Measurement of WBC counts in serum to evaluate the potential effect of preclinical regimens on the immune system and tissue homeostasis of C57BL/6 mice. (i) Percentage of WBC constituents. EO (eosinophil), BASO (basophil), MONO (monocyte), LYMPH (lymphocyte) and NEUT (neutrophil) cell subpopulations were determined separately to categorize these subpopulations. (j-l) Measurement of blood cell components and circulating factors, including RBC count (j, /L), HGB amount (k, g/L) and PLT number (l, /L) in the peripheral blood of animals. (m-p) Routine tests of biochemical parameters. Terminal bleeds were taken from retro-orbital of animals at the end of therapeutic regimens, serum concentrations of creatinine (m, mg/dl), urea (n, mg/dl), ALP (o, U/L) and ALT (p, U/L)) were determined for *in vivo* toxicity appraisal. H&E, hematoxylin and eosin. IHC, immunohistochemistry. IF, immunofluorescence. WBC, white blood cell. RBC, red blood cell. HGB, hemoglobin. PLT, platelets. ALP, alkaline phosphatase. ALT, alanine aminotransferase. Data in b, c, e-p are shown as mean ± SD and representative of 3 independent biological replicates. $P$ values were calculated by Student's $t$ tests. ^, $P > 0.05$; ^, $P > 0.05$; *, $P < 0.05$; **, $P < 0.01$; ***, $P < 0.001$; ****, $P < 0.0001$.
